# Mct11 deficiency alters hepatic glucose metabolism and energy homeostasis

**DOI:** 10.1101/2021.09.08.459307

**Authors:** Alina Ainbinder, Liping Zhao, Patricia Glover, Karen Gelinas-Roa, Victor Rusu, Alycen Harney, Eitan Hoch, Amy A. Deik, Kerry A. Pierce, Kevin Bullock, Courtney Dennis, Sarah Jeanfavre, Jesse Krejci, Jinyoung Choi, Anthony N. Hollenberg, Federico Centeno-Cruz, Francisco Barajas-Olmos, Carlos Zerrweck, Lorena Orozco, Clary B. Clish, Eric S. Lander, Jose C. Florez, Suzanne B. R. Jacobs

## Abstract

Genetic variation at the *SLC16A11* locus contributes to the disproportionate impact of type 2 diabetes (T2D) on Latino populations. We recently demonstrated that T2D risk variants reduce *SLC16A11* liver expression and function of MCT11, the monocarboxylate transporter encoded by the *SLC16A11* gene. Here, we show that *SLC16A11* expression within the liver is primarily localized to the low oxygen pericentral region, and that T2D risk variants disrupt oxygen-regulated *SLC16A11* expression in human hepatocytes. Under physiologic oxygen conditions, MCT11 deficiency alters hepatocyte glucose metabolism, resulting in elevated intracellular lactate and a metabolic shift toward triacylglycerol accumulation. We also demonstrate an impact of Mct11 deficiency on glucose and lipid metabolism in *Slc16a11* knockout mice, which display physiological changes that are observed in individuals with T2D. Our findings provide mechanistic insight into how *SLC16A11* disruption impacts hepatic energy metabolism and T2D risk, and highlight MCT11-mediated regulation of lactate levels as a potential therapeutic target.

## INTRODUCTION

Type 2 diabetes (T2D) is a growing public health crisis that currently affects over 463 million people worldwide and is a leading cause of morbidity and mortality, increasing risk for cardiovascular disease, non-alcoholic fatty liver disease (NAFLD), kidney disease, blindness, and stroke (Portillo-Sanchez et al., 2015; Zheng et al., 2018). T2D is a complex and heterogeneous disorder defined by a common endpoint—hyperglycemia—that ultimately results from a combination of impaired insulin response in peripheral tissues such as liver, adipose, and skeletal muscle, and a decline in pancreatic beta-cell function leading to loss of insulin secretion.

While T2D is influenced by environmental factors, it is also a highly heritable disorder for which over 700 distinct genetic associations across ancestries have been implicated in disease risk (Mahajan et al., 2018; Spracklen et al., 2020; Vujkovic et al., 2020). This genetic foundation of T2D provides an opportunity to elucidate causal disease mechanisms—through identification of the molecular, cellular, and physiological processes impacted by T2D-associated variants—and improve patient care by identifying individuals at risk of developing T2D, guiding more effective treatment approaches, and highlighting new target genes and pathways for drug development (Aquilante, 2010; Gloyn et al., 2004; Inzucchi, 2002; Pearson et al., 2003; Plenge et al., 2013; Shepherd et al., 2009).

T2D-associated variants at the *SLC16A11* locus, located at *17p13*, are among those conferring the highest risk of T2D among common variants reported to date, with each risk allele increasing T2D risk by ∼25% (SIGMA T2D Consortium et al., 2014). T2D risk at this locus is conferred by a haplotype of 18 tightly linked variants, including five coding variants in the *SLC16A11* gene and three non-coding variants near the *SLC16A11* transcriptional start site (Rusu et al., 2017). With an allele frequency of ∼30-50% in populations of Native American ancestry and <2% in populations of European ancestry, this haplotype may account for ∼20% of the increased T2D prevalence in Mexico and U.S. Hispanic populations (SIGMA T2D Consortium et al., 2014).

In previous mechanistic studies, we established that genetic variants on the *SLC16A11* T2D risk haplotype have two distinct effects, both of which lead to reduced function of MCT11, the proton-coupled monocarboxylate transporter (MCT) encoded by the *SLC16A11* gene: regulatory variants decrease *SLC16A11* expression in liver, and coding variants reduce levels of the transporter at the cell surface (Rusu et al., 2017). Furthermore, disruption of *SLC16A11* expression in primary human hepatocytes (PHH) leads to changes in fatty acid and lipid metabolism—in particular, increased diacylglycerols (DAGs) and triacylglycerols (TAGs)—that are also observed in the pathophysiology of insulin resistance and T2D (Rusu et al., 2017). These results implicate reduced MCT11 function in liver as a causal risk factor for T2D and raise the hypothesis that treatment paradigms that increase MCT11 function, or target the metabolic pathways through which MCT11 acts, could benefit individuals with T2D.

Though genetic and experimental data clearly implicate MCT11 function in liver as impacting T2D risk, its specific role in hepatic metabolism and how it impacts glycemic physiology are not clear. MCT11 belongs to a subgroup of the MCT family that transport simple monocarboxylates through a proton-coupled mechanism (Rusu et al., 2017). While MCT11 has been shown experimentally to transport pyruvate (Rusu et al., 2017), closely related family members also transport other monocarboxylates, such as lactate and ketone bodies (Halestrap, 2013). Therefore, it is likely that MCT11 also transports additional monocarboxylates which, along with pyruvate, have a central role in energy metabolism in the liver. Furthermore, since other MCT family members with overlapping substrates are also expressed in the liver (SIGMA T2D Consortium et al., 2014), it is not known how MCT11 deficiency in the liver distinctly alters hepatic metabolism.

Metabolic function within liver lobules is distributed across zones of hepatocytes that are induced by gradients of oxygen, nutrients, and hormones (Bhatia et al., 1996; Jungermann and Kietzmann, 1997; Trefts et al., 2017). These local microenvironments result in subpopulations of hepatocytes that express different enzymes and solute carriers, and therefore specialize in different metabolic processes (Berndt et al., 2021; Kietzmann, 2017; Tachikawa et al., 2018). Whereas hepatocytes located near the oxygen- and nutrient-rich hepatic artery and portal vein (periportal) have a predominant role in gluconeogenesis, fatty acid beta-oxidation, and oxidative phosphorylation, hepatocytes in the relatively oxygen- and nutrient-depleted region near the central vein (pericentral) are the primary site for glucose uptake, glycolysis, lipogenesis, and TAG synthesis (Bazotte et al., 2014; Kietzmann, 2017; Trefts et al., 2017). T2D-associated insulin resistance in the liver is characterized by increased hepatic glucose output, impaired mitochondrial function, accumulation of free fatty acids (FFAs) and lipids, and increases in oxidative stress and inflammation (Boden and Shulman, 2002; DeFronzo et al., 2015; Perry et al., 2014). Thus, metabolic dysfunctions in both periportal and pericentral regions of the liver are implicated in T2D, and understanding the spatial distribution of MCT11 within the liver may help clarify its role in hepatic metabolism and T2D risk.

Here, we set out to elucidate the specific role of MCT11 in hepatic metabolism and physiology. We identify the low oxygen, pericentral environment as an essential context for understanding T2D risk haplotype effects on *SLC16A11* expression and MCT11 function in liver. We establish a crucial role for MCT11 in regulating lactate levels in hepatocytes exposed to pericentral oxygen tension, and identify lactate as a potential circulating biomarker of Mct11 function in an *Slc16a11* knockout mouse model. We further demonstrate that Mct11 deficiency leads to changes in energy metabolism and physiology observed in individuals with T2D, including accumulation of hepatic and circulating TAGs. Together, our findings suggest that T2D risk haplotype disruption of MCT11 increases disease risk due to a shift in energy homeostasis that emulates nutrient excess and increases metabolic stress, which can ultimately lead to inflammation, liver damage, insulin resistance, and T2D.

## RESULTS

### *SLC16A11* expression in the liver is enriched in the pericentral hepatocyte population

To gain insight into how genetic disruption of *SLC16A11* alters hepatic metabolism despite low expression levels and overlapping function with other SLC16 family members (Rusu et al., 2017; SIGMA T2D Consortium et al., 2014), we investigated where *SLC16A11* is expressed within the liver, a heterogeneous tissue in which metabolic function is distributed between periportal and pericentral zones of hepatocytes (Trefts et al., 2017). We first analyzed single-cell RNA sequencing (RNA-seq) data from mouse liver (Halpern et al., 2017), and found that *SLC16A11* shows a zonal expression profile, with significantly higher expression in pericentral-zone hepatocytes compared to periportal-zone hepatocytes. *SLC16A11* expression was not detected in other liver cell types, including endothelial or Kupffer cells (Figures 1A-B) (Halpern et al., 2018). In contrast, closely related SLC16 family members that are likely to have overlapping transport function (Halestrap, 2013; Rusu et al., 2017), such as *SLC16A1* and *SLC16A13*, are more broadly expressed across liver cell types and do not display zonal expression profiles within hepatocytes (Figures 1A-B). Consistent with these mouse data, human *SLC16A11* also shows a pericentral profile, as measured through laser-capture microdissection (LCM) RNA-seq of liver zones (Figure 1C) (McEnerney et al., 2017). The discovery that *SLC16A11* is predominantly expressed in pericentral hepatocytes, which represent a fraction of cells within the liver, provides an explanation for the low *SLC16A11* expression levels detected in bulk liver tissue analyses (SIGMA T2D Consortium et al., 2014), and highlights one way in which *SLC16A11* is distinct from closely related family members.

**Figure 1.**
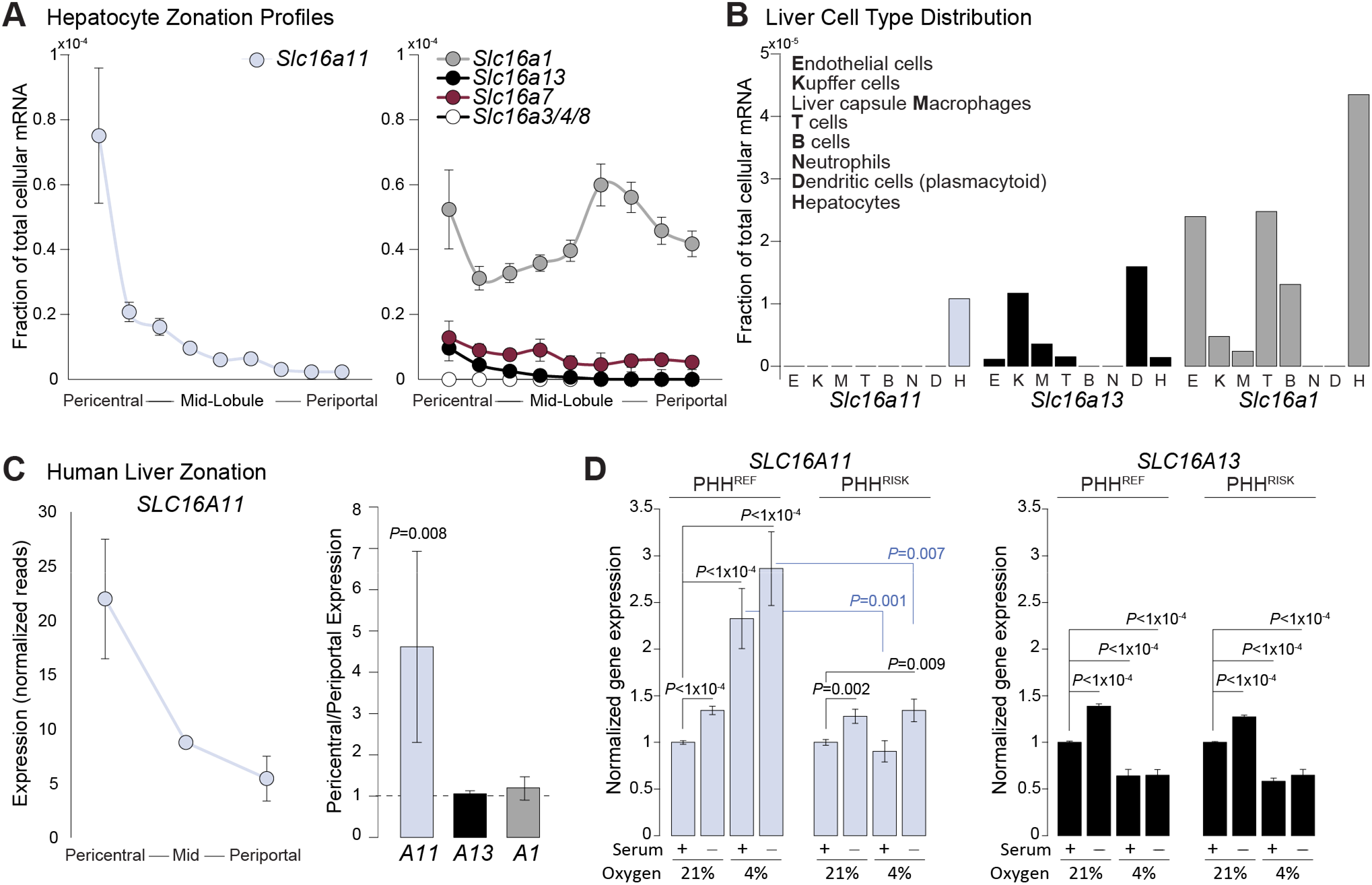
T2D risk variants disrupt *SLC16A11* regulation by pericentral oxygen tension. (A) Zonation profile of mouse *Slc16a11* expression measured through single-cell RNA sequencing (Halpern et al., 2017). (B) Cell type distribution of *Slc16a11, Slc16a13*, and *Slc16a1* expression within the liver (Halpern et al., 2018). (C) Expression of *SLC16A11* across human liver zones quantified through laser-capture microdissection RNA-sequencing (LCM-RNAseq) (left), and relative pericentral-to-periportal expression of *SLC16A11*, *SLC16A13*, and *SLC16A1* in LCM-RNAseq data (right) (McEnerney et al., 2017). (D) *SLC16A11* and *SLC16A13* expression in primary human hepatocytes homozygous for the *SLC16A11* reference (PHH^REF^) or T2D risk (PHH^RISK^) haplotypes, under standard 21% oxygen and physiologically relevant (4%) pericentral oxygen tension, in the presence and absence of serum. Bar plots depict relative gene expression ± SEM normalized to the 21% with serum condition within each genotype. Significant *P* values are indicated for effects of serum withdrawal and/or 4% oxygen relative to 21% oxygen with serum conditions within PHH for each genotype and comparisons between PHH^REF^ and PHH^RISK^ effects for each environmental exposure. *SLC16A11* expression: n=4 PHH^REF^ donors and n=2 PHH^RISK^ donors; *SLC16A13* expression: n=2 PHH^REF^ donors and n=2 PHH^RISK^ donors.

### T2D risk variants disrupt *SLC16A11* regulation by physiologic pericentral oxygen tension

To facilitate mechanistic inquiry, we next sought to determine if we could replicate the zonal regulation of *SLC16A11* in primary human hepatocytes (PHH) *in vitro*. Standard cell culture conditions are more akin to the nutrient- and oxygen-rich periportal region; therefore, to model the pericentral region, we cultured PHH under physiologic pericentral oxygen tension (4%) and low-nutrient conditions (serum-free media) for 48 hours (Jungermann and Kietzmann, 1997). *SLC16A11* expression is induced ∼30% in the absence of serum and increased 2.5-fold by physiologic 4% oxygen tension in PHH homozygous for the *SLC16A11* reference, non-risk haplotype (PHH^REF^) (*P*<1×10^−4^, Figure 1D).

We then examined the effect of the T2D-associated haplotype on expression in this model system. Notably, in PHH homozygous for the T2D risk haplotype (PHH^RISK^), low oxygen fails to increase *SLC16A11* expression (Figure 1D). The T2D risk haplotype had no effect on *SLC16A11* levels induced by the absence of serum-containing nutrients, hormones, and growth factors. In contrast to the effect of oxygen on *SLC16A11*, hepatocyte expression of *SLC16A13*, the adjacent gene and a closely related family member, is lower at 4% oxygen (*P*<1×10^−4^) and not impacted by the T2D risk haplotype (Figure 1D). Taken together, our data suggest the previously identified eQTL between the T2D risk haplotype and decreased *SLC16A11* in liver (Rusu et al., 2017) may be primarily attributable to loss of pericentral *SLC16A11* expression. Furthermore, these data establish a link between oxygen-induced *SLC16A11* expression in hepatocytes and T2D risk, pointing toward a key a role for MCT11 action in pericentral hepatocyte metabolism.

### MCT11 regulates lactate levels in hepatocytes exposed to pericentral oxygen tension

Having established zone- and T2D risk haplotype-dependent expression of *SLC16A11*, we sought to understand the effects of this discovery on cellular metabolism. The low oxygen microenvironment where *SLC16A11*-expressing pericentral hepatocytes reside is key to the metabolic function of these cells—which are highly dependent on glycolysis for energy production (Kietzmann, 2017; Trefts et al., 2017). Moreover, MCT11 belongs to a category of MCTs that transport pyruvate and lactate—the end products of glycolysis and fuel for the TCA cycle. We therefore used comprehensive metabolite profiling to assess the impact of *SLC16A11* loss on metabolic processes in hepatocytes in response to glucose stimulation at physiologically relevant 4% oxygen. Using pooled siRNAs with proven specificity (Rusu et al., 2017), we achieve ∼75% knock down of *SLC16A11* expression compared to non-targeting siRNA-treated PHH at the corresponding oxygen tension (Figure 2A), a degree of perturbation that is comparable to the T2D risk variant haplotype effect on MCT11 in humans (Rusu et al., 2017). To ascertain MCT11 effects on both primary and downstream glucose metabolic pathways, we exposed control and *SLC16A11* knockdown PHH to the different oxygen tensions for 48 hours, and generated metabolite profiles at baseline and at multiple time points following stimulation with fresh glucose-containing media (Figure S1A).

**Figure 2.**
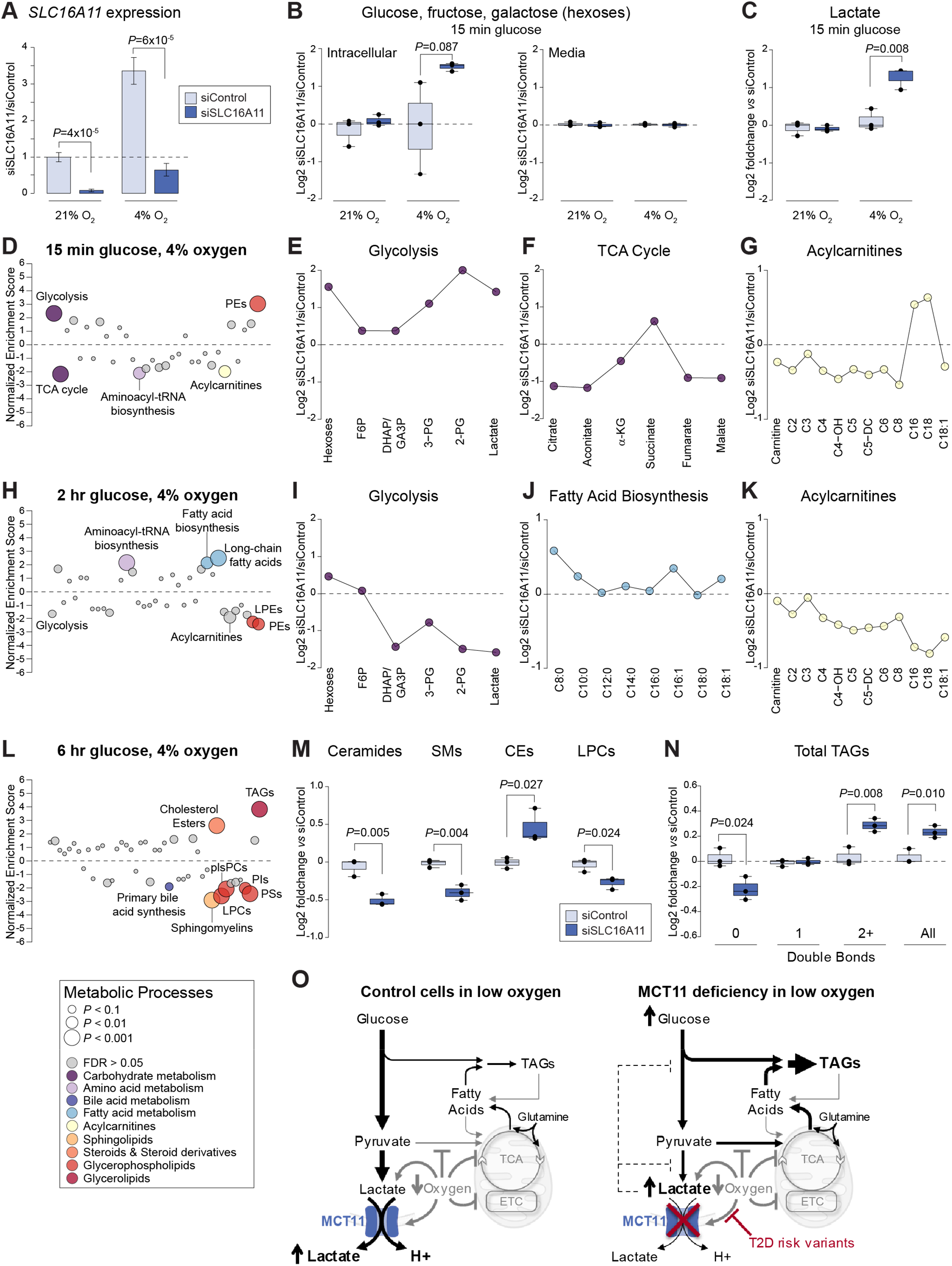
MCT11 deficiency alters lactate export from PHH at physiological oxygen. (A) *SLC16A11* expression in PHH treated with siRNAs targeting *SLC16A11* (siSLC16A11) or control siRNAs (siControl) in cells cultured at 21% or 4% oxygen tension for 48 h. Bar plots depict relative gene expression ± SEM using TBP for normalization. (B) Relative levels of intracellular and extracellular (media) hexose sugars (glucose, fructose, galactose) in control and *SLC16A11* knockdown PHH cultured at 21% and 4% oxygen and after 15 min stimulation with fresh glucose-containing media. Boxplots depict log2 foldchange relative to control knockdown PHH. (C) Intracellular lactate levels in control and *SLC16A11* knockdown PHH cultured at 21% and 4% oxygen after 15 min exposure to fresh glucose. Boxplots depict log2 foldchange relative to control knockdown PHH. (D-G) Intracellular metabolic pathway changes following 15 min stimulation with fresh glucose-containing media in *SLC16A11* knockdown PHH cultured at 4% oxygen. (D) Pathway enrichment analysis. Each dot represents a different metabolic pathway or metabolite class. *P* values are indicated by dot size. Significantly altered pathways and metabolite classes (false discovery rate [FDR] < 0.05) are labeled and colored according to KEGG metabolic process or metabolite class (see legend below Figure 2L), with non-significant pathways shown in gray. PEs, phosphatidylethanolamines. (E-G) Line plots showing log2 fold changes in individual metabolites in (E) glycolysis, (F) the TCA cycle, and (G) acylcarnitines. (H-K) Intracellular metabolic pathway changes following 2 hr stimulation with fresh glucose-containing media in *SLC16A11* knockdown PHH cultured at 4% oxygen. (H) Pathway enrichment analysis as described above. (I-K) Line plots showing log2 fold changes in individual metabolites in (I) glycolysis, (J) fatty acid biosynthesis, and (K) acylcarnitines. LPEs, lysophosphatidylethanolamines. (L-N) Intracellular metabolic pathway changes following 6 hr stimulation with fresh glucose-containing media in *SLC16A11* knockdown PHH cultured at 4% oxygen. (H) Pathway enrichment analysis as described above. LPCs, lysophosphatidylcholines; plsPCs, phosphatidylcholine plasmalogens; PIs, phosphatidylinositols; PSs, phosphatidylserines. (M-N) Total levels of (M) ceramides, sphingomyelins (SMs), cholesterol esters (CEs), and LPCs, and (N) triacylglycerols (TAGs), stratified by saturation and combined. Double bonds: 0=saturated, 1=monounsaturated, 2+=polyunsaturated. Boxplots depict log2 foldchange relative to control knockdown PHH. (O) Model depicting the effect of 4% oxygen on normal glucose metabolism (left) and the effect of MCT11 deficiency on glucose metabolism in this environment (right). TCA, tricarboxylic acid cycle; ETC, electron transport chain. See also Figure S1 and Tables S1 and S2.

To confirm expected environment-induced metabolic changes in our cell model, we first evaluated the effect of 4% oxygen compared to 21% oxygen in control PHH at baseline. Control PHH residing at 4% oxygen for 48 hours have significantly higher accumulation of both intracellular and extracellular lactate, indicative of the increased glycolytic state of these cells (Figure S1B). Similarly, cells in low oxygen tension display metabolic changes signifying decreased fatty acid oxidation (reduced acylcarnitines), and increases in ketogenesis (elevated α-/β-hydroxybutyrate), bile acid synthesis (elevated intracellular bile acid levels), and lipogenesis (increased TAGs) (Figures S1C-F and Table S1). These pathway changes are consistent with metabolic functions associated with the pericentral zone (Trefts et al., 2017).

We tested whether MCT11 deficiency affects pathways involved in central carbon metabolism in PHH after acute exposure to fresh glucose-containing media. In cells cultured at 4% oxygen and stimulated with glucose for 15 minutes, levels of intracellular hexose sugars (fructose/glucose/galactose) are ∼3-fold higher in *SLC16A11* knockdown cells compared to control cells despite comparable levels in the media (Figure 2B). At this time point, the metabolite most significantly affected by MCT11 deficiency is intracellular lactate, which is increased ∼2.7-fold (*P*=0.008; Figure 2C and Table S1). Levels of α- and β-hydroxybutyrate, which are other monocarboxylates transported by some MCT family members, are also increased about 1.6-fold (*P*=0.04) (Table S1).

We analyzed the overall effect of MCT11 deficiency on metabolic pathways and processes using enrichment analyses that examined changes in KEGG metabolic pathways and classes of functionally and structurally related metabolites. These pathway analyses identified increased levels of glycolytic metabolites (*P_GSEA_*<0.001), decreased levels of TCA cycle metabolites (*P_GSEA_*<0.001), and decreased fatty acid oxidation intermediates (acylcarnitines; *P_GSEA_*=0.008) in MCT11-deficient PHH 15 min after glucose stimulation (Figures 2D-G and Table S2). Notably, the accumulation of lactate and other glycolytic metabolites and the decrease in TCA cycle metabolites were only observed in cells cultured at low oxygen (Figures 2C and S1G-I)—which promotes production of lactate and inhibits pyruvate entry into the TCA cycle (Cui et al., 2017; Kim et al., 2006; Papandreou et al., 2006). These data underscore the importance of the low oxygen environment to MCT11 action, suggest MCT11 is a key regulator of lactate levels in pericentral hepatocytes, and reveal an acute effect of MCT11 deficiency on central carbon metabolism manifested as the intracellular accumulation of lactate and other glycolytic metabolites.

### MCT11 deficiency leads to triacylglycerol accumulation in hepatocytes

To evaluate downstream consequences of MCT11 deficiency on glucose-stimulated fatty acid and lipid levels at 4% oxygen, we profiled control and *SLC16A11* knockdown PHH at 2 and 6 hours after glucose stimulation. Two hours after stimulation with fresh glucose-containing media, MCT11-deficient PHH have higher levels of free fatty acids (FFA) (fatty acid biosynthesis, *P_GSEA_*=0.004 and long-chain fatty acids (*P_GSEA_*<0.001) and decreased fatty acid oxidation (acylcarnitines, *P_GSEA_*=0.009) (Figures 2H-K and Table S2). Though elevated at the 15 minute time point, levels of metabolites in the energy-producing, NAD+-dependent phase of glycolysis are significantly lower in *SLC16A11* knockdown PHH 2 hours after fresh media stimulation [Dihydroxyacetone phosphate/glyceraldehyde 3-phosphate (DHAP/GA3P) *P*=0.0009; 3-phosphoglycerate (3-PG) *P*=0.01; 2-phospoglycerate (2-PG) *P*=0.02] (Figure 2I and Table S1), potentially due to feedback inhibition by lactate and/or decreased NAD+ availability (Benjamin et al., 2018; Costa Leite et al., 2007; Leite et al., 2011).

Six hours after stimulation with fresh glucose-containing media, MCT11-deficiency has a substantial impact on hepatocyte lipid composition, resulting in decreased levels of sphingolipids (sphingomyelins (SMs) *P_TOTAL_*=0.004 and ceramides *P_TOTAL_*=0.005) and phospholipids and increased levels of intracellular cholesterol esters (CEs; *P_TOTAL_*=0.02) and TAGs (*P_TOTAL_*=0.01) (Figures 2L-N and Table S1). The increase in total TAGs is driven by an increase in polyunsaturated TAGs (*P_TOTAL_*=0.008) (Figure 2N), potentially due to impacts on redox state and consistent with what was observed in human carriers of the T2D risk haplotype (Kim et al., 2019).

Overall, our *in vitro* data demonstrate that MCT11 has a critical role in regulating cellular lactate levels in the low oxygen pericentral liver microenvironment, and suggest that MCT11 deficiency causes a shift in central carbon metabolism that promotes TAGs accumulation (Figure 2O), which may herald hepatic dysfunction, IR, and T2D.

### *Slc16a11*-deficient mice are viable and have normal body weight and body composition

To investigate the contribution of *Slc16a11* to metabolic physiology, we generated a knockout mouse model using CRISPR/Cas9 technology to introduce a 19 bp deletion in the second coding exon of the mouse gene, creating a frameshift at amino acid 94, and early termination (p.Gly94SerfsTer34) (Figures 3A-B). Two chimeric founder mice with the same mutation (*Slc16a11^del19^*) were bred to C57BL/6J mice to create a germline knockout model, and remove potential non-linked, off-target effects. Expression analyses by droplet digital PCR (ddPCR) in adult liver show an approximate 70% reduction in *Slc16a11* expression, and confirm that transcript levels of other Slc16 family members and nearby genes (*Slc16a13, Bc6lb*) are not affected by the genetic perturbation (Figure 3C). Examination of *Slc16a11* levels in other T2D-relevant metabolic tissues detected low expression in subcutaneous adipose, pancreas, and thyroid and undetectable levels in skeletal muscle (Figure S2A).

**Figure 3.**
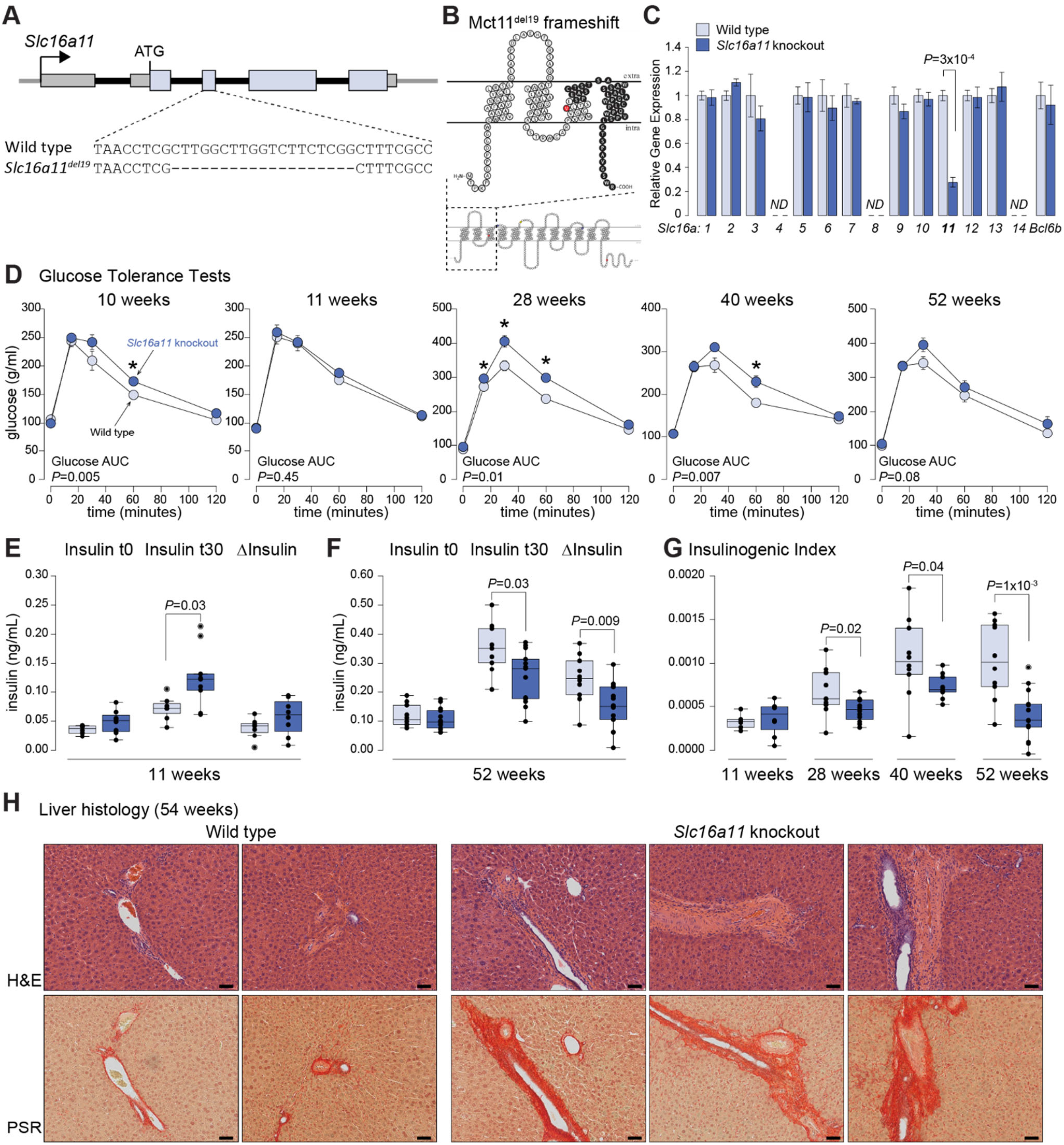
Effect of *Slc16a11* deficiency on glycemic physiology and liver health. (A) Depiction of mouse *Slc16a11* locus showing 19 bp deletion introduced into *Slc16a11^del19^* knockout mouse model. (B) Predicted truncation and membrane topology of mouse Mct11^del19^. Black circles represent frame-shifted region of protein. Red circle indicates location of V89, the amino acid analogous to human V113, which is altered by a variant on the T2D risk haplotype. Wild-type mouse Mct11 is shown below. (C) Expression of Slc16 family members and *Bcl6b* (a nearby gene) in livers from adult wild-type (n=6) and *Slc16a11* homozygous knockout (n=4) mice. Bar plots depict gene expression relative to wild-type mice ± SEM using TBP for normalization. *ND*=not detected. (D) Intraperitoneal glucose tolerance tests (IPGTT) from cohort A at 10, 11, 28, 40, and 52 weeks of age. * *P* < 0.05 for an individual time point during IPGTT. n=11-12 wild-type and 14 *Slc16a11*-knockout mice. (E-F) Insulin measured at fasting (t0) and 30 min after glucose administration (t30) during IPGTTs in (E) 11 week old and (F) 52 week old mice. ΔInsulin = Insulin (t30-t0). (G) Insulinogenic index [Insulin (t30-t0)/Glucose (t30-t0)] calculated during IPGTTs in mice at 11, 28, 40, and 52 weeks of age. (H) H&E and picrosirius red (PSR, collagen) staining of liver sections from 54 week old fasting wild-type and *Slc16a11* knockout mice. Images shown are from different animals and were taken using a 20x (200x, scale bar = 50 μM) objective. See also Figure S2.

To obtain cohorts of wild-type and knockout littermates for metabolic phenotyping, we intercrossed heterozygous *Slc16a11^del19^* mice. The number of homozygous *Slc16a11*-knockout mice present at weaning is consistent with Mendelian ratios (wild-type: *n*=841, 24.8%; heterozygous: *n*=1678, 49.6%; homozygous knockout: *n*=863, 25.5%), demonstrating *Slc16a11*-knockout mice are viable. In addition, *Slc16a11-*knockout animals are fertile and overtly indistinguishable in appearance from their wild-type littermates. We observed no differences in body weight or body composition over the course of 12 months in mice fed a normal chow diet (Figures S2B and S2C).

### *Slc16a11* is not essential for maintaining glucose homeostasis in mice

To test the effect of *Slc16a11* loss on glycemic physiology *in vivo*, we carried out longitudinal studies of cohorts consisting of sex- and litter-matched wild-type and *Slc16a11-*knockout animals. For each of five cohorts, we conducted multiple intraperitoneal glucose tolerance tests (IPGTT) between 2 to 6+ months of age, and observed varying effects of *Slc16a11* deficiency on glucose tolerance. Specifically, *Slc16a11*-knockout mice displayed reduced glucose tolerance compared to their wild-type littermates in three cohorts, and no glycemic differences by genotype were observed in the other two cohorts (Figures 3D and S2D-E). In *Slc16a11*-knockout mice with significant reduced glucose tolerance, we sought to determine if *Slc16a11* deficiency was associated with additional, long-term changes in glycemic physiology that might provide mechanistic insight. We measured insulin response during the IPGTTs and found that the elevated glucose levels observed in *Slc16a11*-knockout mice is not due to a primary deficiency in insulin secretion, as young, adult animals are able to compensate and manage the higher glucose load through increased insulin secretion (Figure 3E, *11 weeks*). However, these *Slc16a11*-knockout mice lost this adaptive insulin response as they aged (Figures 3F-G, *28-52 weeks*), suggesting they may follow a pattern of beta-cell burnout similar to what is observed during T2D progression in humans (Chen et al., 2017; Kahn et al., 1993). We did not detect differences in liver enzyme levels (serum ALT) or in thyroid axis physiology (plasma T4, TSH), which we evaluated due to the known expression of *Slc16a11* in thyroid tissue (SIGMA T2D Consortium et al., 2014) (Figures S2F-G). Examination of histological features in 54-week old wild-type and *Slc16a11*-knockout liver revealed the presence of microsteatosis and inflammation in both genotypes; however, we observed increased immune cell infiltration in livers from *Slc16a11* deficient mice (Figure S2H). Notably, four out of five *Slc16a11*-knockout livers harbor fibrotic regions with excessive connective tissue on compared to what is observed in their wild-type littermates (Figure 3H). We identified no obvious histological differences in skeletal muscle, adipose, or pancreas (Figure S2I). Although these data indicate *Slc16a11* is not required to maintain glucose homeostasis in mice, they point to a potential impact of *Slc16a11* deficiency on glucose metabolism in insulin-sensitive peripheral tissues, and demonstrate a role in maintaining liver health.

### *Slc16a11* deficiency induces a shift toward TAG storage and metabolic stress *in vivo*

To gain further insight into underlying metabolic changes that may contribute to altered pathophysiology over time, we examined *in vivo* molecular and cellular changes caused by *Slc16a11* deficiency in 17-week old adult mice after overnight fasting and in response to a 30 minute glucose challenge (Figure S3A). At the time of sample collection, these mice did not exhibit differences in blood glucose or insulin measures (Figure S3B); therefore, metabolic phenotypes represent primary genetic effects on metabolism rather than a secondary consequence of a hyperglycemic state. We measured biochemical consequences of *Slc16a11* loss through comprehensive metabolite profiling of ∼600 known polar, fatty acid, and lipid metabolites in liver and plasma from wild-type and *Slc16a11-*knockout mice.

In fasting animals, individual metabolite and pathway analyses highlight changes in fatty acid and lipid metabolism akin to what was observed *in vitro* (Figures 4A-B, and Tables S3-S4). These data show a significant enrichment for elevated liver TAGs in *Slc16a11*-deficient mice compared to wild-type littermates, leading to a cumulative ∼15% higher total liver TAG content (*P_TOTAL_*=0.036) that is driven by higher levels of unsaturated TAGs across all acyl chain lengths (Figures 4A, 4C-D, and S3C). This increase in liver TAGs is accompanied by decreases in fasting FFAs (long-chain fatty acids *P_GSEA_*=0.002), sphingolipids (ceramides *P_TOTAL_*=9×10^−4^ and SMs *P_TOTAL_*=8×10^−4^), and phospholipids [lysophosphatidylethanolamines (LPEs) *P_TOTAL_*=0.006 and phosphatidylcholines (PCs) *P_TOTAL_*=0.008] (Figures 4A and 4E-F). Similar changes in fatty acid and lipid metabolism were detected in plasma, with fasting *Slc16a11*-deficient mice showing enrichment for decrease in circulating sphingolipids (*P_GSEA_*<0.001) and higher circulating medium- and long-chain acylcarnitines (*P_GSEA_*<0.001) and TAGs (*P_GSEA_*<0.001) (Figure 4B and S3D). The plasma TAG profile specifically shows increased levels of TAGs containing fewer acyl chain carbons (Figure S3C), similar to the circulating TAG profile that is associated with insulin resistance and higher risk of future T2D in humans (Rhee et al., 2011). Beyond demonstrating that Mct11 deficiency *in vivo* induces multiple metabolic phenotypes that are observed in humans with or at risk of developing T2D (Adams et al., 2009; Mai et al., 2013; Mihalik et al., 2010; Seymour and Byrne, 1993; Stern and Haffner, 1991), these data are consistent with the cellular findings in PHH, and reveal that Mct11 deficiency induces a metabolic shift towards glycerolipid energy storage under fasting conditions that are typically expected to stimulate utilization of energy stores.

**Figure 4.**
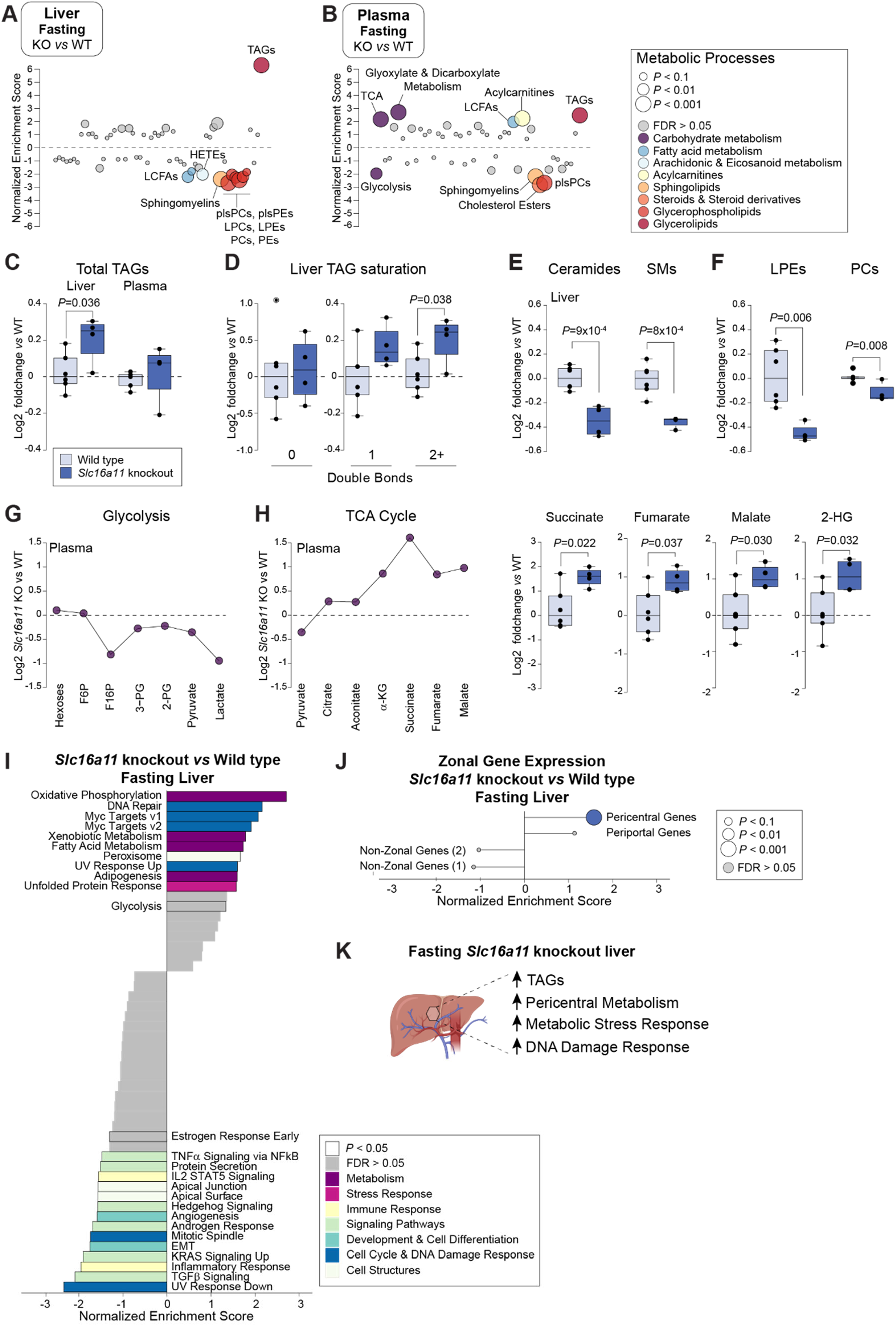
Fasting *Slc16a11* deficient mice have increased metabolic stress and hepatic triacylglycerol storage. (A-B) Enrichment analysis of metabolic pathway changes in (A) liver and (B) plasma from fasting *Slc16a11*-knockout (KO) mice compared to wild-type (WT) littermates. LCFAs, long-chain fatty acids; HETEs, hydroxyleicosatetraenoic acids; plsPEs, phophatidylenthalomine plasmalogens, LPCs, lysophosphatidylcholines; PIs, phosphatidylinositols; PSs, phosphatidylserines. (C-F) Total levels of (C) liver and plasma TAGS, (D) liver TAGs stratified by saturation, (E) liver sphingolipids (ceramides and SMs), and (F) liver phospholipid (LPCs and phosphatidylcholines (PCs)) from fasting wild-type and *Slc16a11*-knockout mice. Double bonds: 0=saturated, 1=monounsaturated, 2+=polyunsaturated. Boxplots depict log2 foldchange relative to wild-type mice. (G-H) Individual metabolite changes in (G) glycolysis and (H) the TCA cycle detected in plasma from *Slc16a11*-knockout mice compared to wild-type littermates. Line plots and boxplots depict log2 foldchange relative to wild-type mice for individual metabolites. 2-HG; 2-hydroxyglutarate. (I) Gene set enrichment analysis (GSEA) of liver transcriptomic profiles from fasting *Slc16a11*-knockout mice compared to wild-type littermates. Normalized enrichment score (NES) indicates increased (positive) or decreased (negative) expression in *Slc16a11* knockout. Each bar represents a different pathway from the Hallmark pathways collection (Liberzon et al., 2015). Pathways with *P* < 0.05 are labeled and outlined. Colors represent different categories of pathways. Non-significant pathways with (false discovery rate [FDR] > 0.05) are shown in gray. (J) GSEA of expression changes in genes with pericentral and periportal zonation profiles in fasting *Slc16a11*-knockout versus wild type livers. (K) Summary of metabolic effects of Mct11 deficiency observed at fasting. See also Figures S3 and S4 and Tables S4 and S5.

In addition to effects on lipid metabolism, fasting *Slc16a11*-deficient mice have altered levels of metabolites involved in pathways of central carbon metabolism, including lower levels of circulating glycolytic metabolites (*P_GSEA_*=0.001) and elevated levels of circulating TCA cycle metabolites (*P_GSEA_*<0.001), in particular aconitate (*P*=0.047), succinate (*P*=0.022), fumarate (*P*=0.037), and malate (*P*=0.030) as well as 2-hydroxyglutarate (*P*=0.032), an α-ketoglutarate-derived metabolite (Figures 4B and 4G-H and Table S3). In liver, levels of these TCA cycle metabolites are similar in *Slc16a11*-deficient and wild-type mice (Figure S3E and Table S3); however, *Slc16a11*-deficient mice have a lower glutamine-to-glutamate ratio (Figure S3F), a prognostic marker for T2D risk that may indicate altered utilization of glutamine to fuel the TCA cycle (Walford et al., 2016; Yang et al., 2014).

To further characterize these metabolic changes, we assessed levels of relevant transcripts altered by *Slc16a11* deficiency through RNA-sequencing of liver samples from wild-type and knockout mice. Analyses of individual gene differences demonstrate *Slc16a11* is among the most down-regulated genes, and confirm our ddPCR results showing no effect of the 19 bp deletion on expression of other Slc16 family members or nearby genes (Figures S4A-B). We used gene set enrichment analysis (GSEA) to assess genotype-driven differences in hepatic expression of 50 hallmark gene sets that represent well-defined biological processes (Liberzon et al., 2015). Among fasting animals, these analyses revealed that Mct11-deficient mice have higher expression of genes involved in key hepatic metabolic processes, including oxidative phosphorylation, xenobiotic metabolism, and fatty acid metabolism, along with increased expression of stress and damage response genes, such as those involved in the unfolded protein response (UPR), DNA damage response, and DNA repair (Figure 4I and Table S5). Conversely, pathways with reduced expression in livers from fasting *Slc16a11*-deficient mice include those involved in the inflammatory response, TGFβ signaling, epithelial-mesenchymal transition (EMT), and TNFα signaling through NFκB. Taken together, the metabolic and transcriptional changes caused by *Slc16a11* deficiency *in vivo* suggest that the absence of functional Mct11 leads to persistent metabolic stress in the liver, that is present even during periods of fasting.

Because *Slc16a11* is predominantly expressed in pericentral hepatocytes, we investigated whether transcriptional changes in *Slc16a11*-knockout mice primarily affect genes in the pericentral region. Based on spatial expression profiles generated from scRNA-seq analyses of mouse liver (Halpern et al., 2017), we used GSEA to assess expression changes in genes with pericentral zonation profiles, periportal zonation profiles, and control sets of genes that are not zonally distributed. In livers from fasting *Slc16a11*-knockout mice, these analyses revealed an enrichment for increased expression of genes with pericentral zonation profiles and no effect on genes with periportal zonation profiles (Figure 4J). These observations further highlight the importance of *SLC16A11* to maintaining zonal and metabolic homeostasis within the liver (Figure 4K).

### *Slc16a11*-knockout mice have lower circulating lactate after glucose stimulation

Because our data demonstrate that Mct11 impacts the metabolic response to glucose, we also examined metabolite and gene expression differences in wild-type and *Slc16a11*-knockout mice thirty minutes following glucose administration (Figure S3A). Similar to what we observed in PHH *in vitro*, Mct11 deficiency leads to increased hepatic levels of glycolytic metabolites in response to glucose (2-PG, *P*=0.005 and 3-PG, *P*=0.005) (Figure 5A). The most significant individual metabolite difference in the livers of glucose-treated *Slc16a11*-deficient mice is elevated levels of trimethylamine N-oxide (TMAO) (1.5-fold; *P*=0.004) (Figure 5B and Table S3), a metabolic change that activates the UPR and is correlated with hepatic insulin resistance, T2D, and several other cardiometabolic disorders (Chen et al., 2019; Liu et al., 2021; Roy et al., 2020; Subramaniam and Fletcher, 2018).

**Figure 5.**
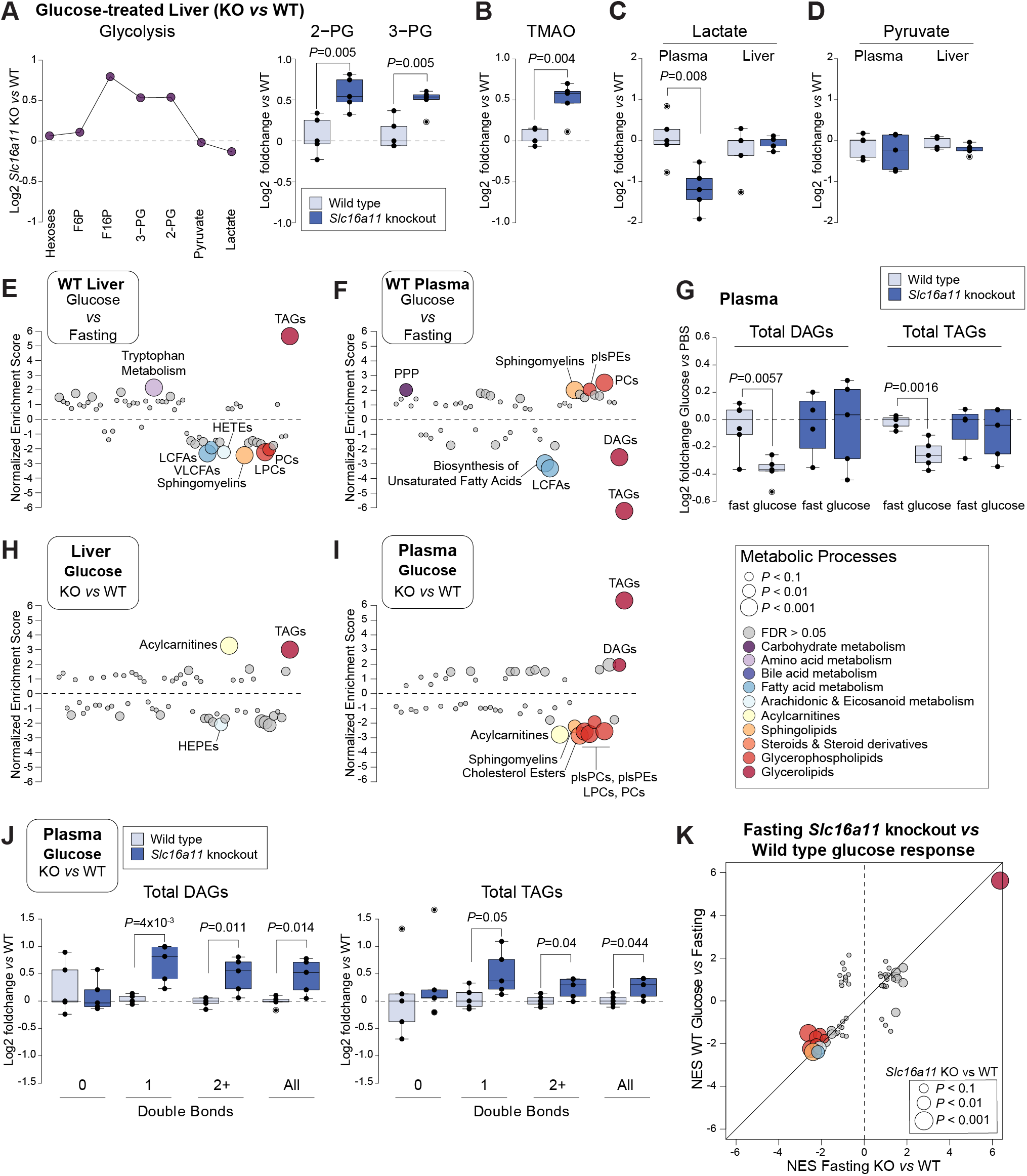
*Slc16a11* deficiency causes a hyperglycemic metabolic state. (A) Individual metabolite changes in glycolysis detected in liver from glucose-treated *Slc16a11*-knockout mice compared to wild-type littermates. Boxplots depict log2 foldchange relative to wild-type mice. See also Table S3. (B) Liver trimethylamine N-oxide (TMAO) levels in wild-type and *Slc16a11*-knockout mice 30 min after glucose administration. Boxplots depict log2 foldchange relative to wild-type mice. (C-D) Plasma and liver (C) lactate and (D) pyruvate levels in wild-type and *Slc16a11*-knockout mice 30 min after glucose administration. Boxplots depict log2 foldchange relative to wild-type mice. (E-F) Enrichment analysis of metabolic pathway changes in (E) liver and (F) plasma in response to glucose in wild-type mice. VLCFAs, very long-chain fatty acids. (G) Total levels of plasma DAGs and TAGs in fasting and glucose-treated wild-type and *Slc16a11*-knockout mice. (H-I) Enrichment analysis of metabolic pathway changes in (H) liver and (I) plasma from glucose-treated *Slc16a11*-knockout mice compared to wild-type littermates. HEPEs, hydroxyeicosapentaenoic acids. (J) Total levels of plasma DAGs and TAGs stratified by saturation and considered all together. Double bonds: 0=saturated, 1=monounsaturated, 2+=polyunsaturated. Boxplots depict log2 foldchange relative to wild-type mice. (K) Scatterplots comparing metabolic pathway changes observed in livers from fasting *Slc16a11*-knockout *vs* wild-type mice with those observed in glucose-treated *vs* fasting wild-type mice. *P* values and FDR are based on fasting *Slc16a11*-knockout *vs* wild-type mice comparison. See also Figure S3 and Tables S3 and S4.

Importantly, we found that plasma lactate levels are ∼55% lower (*P*=0.008) in *Slc16a11*-knockout mice following glucose stimulation (Figure 5C), consistent with the intracellular accumulation of lactate observed in PHH (Figure 2C). This effect on lactate is among the most significant and largest impacts of *Slc16a11* deficiency on circulating metabolites. We did not detect differences in levels of liver lactate (Figure 5C), potentially due to heterogeneity within liver zones, the low proportion of pericentral hepatocytes within the liver, and/or intracellular lactate being metabolized. In addition, we did not detect differences in pyruvate levels in either liver or plasma (Figure 5D). These results suggest a role for MCT11 in glycolysis-driven lactate efflux and indicate that plasma lactate may be a circulating biomarker of MCT11 function.

### *Slc16a11* deficiency causes a metabolic state resembling hyperglycemia in the liver

In addition to comparing wild-type and knockout mice from the same treatment group, we explored the effect of Mct11 on glucose metabolism by comparing how wild-type and *Slc16a11*-deficient mice respond to glucose after fasting (Figure S3A). In wild-type mice, glucose induces metabolic changes consistent with an expected shift towards TAG synthesis and storage (Rui, 2014). This includes an enrichment of elevated TAGs in the liver and lower levels of DAGs (*P_TOTAL_*=0.006) and TAGs (*P_TOTAL_*=0.002) in the circulation (Figures 5E-G). In contrast, *Slc16a11*-deficient mice, which had elevated liver TAGs at fasting, show no further enrichment in liver TAGs following glucose exposure nor do they have the decreases in circulating DAGs and TAGs observed in wild-type mice (Table S4 and Figure 5G). These differences in baseline and glucose response profiles result in less pronounced metabolomic differences between glucose-treated wild-type and *Slc16a11*-deficient livers than what we observed at fasting, but prominent differences in circulating lipid profiles (Figures 5H-I). In plasma, glucose-treated *Slc16a11*-deficient mice have higher total levels of circulating DAGs (1.4-fold, *P_TOTAL_*=0.014) and TAGs (1.2-fold, *P_TOTAL_*=0.044) that are driven by increases in unsaturated lipids (Figure 5J). These distinctions in glucose response are not due to differences in insulin levels (Figure S3B), indicating the observed phenotypes are due to metabolic changes within the liver and potentially other insulin-responsive tissues.

Notably, the consequences of Mct11 deficiency on hepatic metabolism in fasting animals are remarkably similar to the hepatic metabolic pathway changes induced by glucose in wild-type mice (Figure 5K, also compare Figure 5E to 4A). These data suggest that Mct11 deficiency *in vivo* leads to an altered metabolic state whereby the liver is functioning as though exposed to persistent hyperglycemia, thus driving energy storage even under fasting conditions.

### *Slc16a11* deficiency causes metabolic stress and predisposes the liver to an inflammatory response

When we examined hepatic expression differences in glucose-treated mice, we found a dramatic shift from what we observed at fasting. In this context of nutrient exposure, *Slc16a11*-deficient mice have lower expression of metabolic pathway genes and significantly higher expression of inflammatory response genes, including those involved in IFNα, IFNγ, TNFα, NFκB, and cytokine signaling, and genes involved in EMT, an inflammation-induced process implicated in tissue fibrosis (Figure 6A and Table S5) (Kalluri and Weinberg, 2009; Thiery et al., 2009; Wynn and Ramalingam, 2012). These changes in pathway gene expression are accompanied by decreased levels of pericentral genes and increased levels of periportal genes, suggesting an impact on zonal metabolic function (Figures S4C-D).

**Figure 6.**
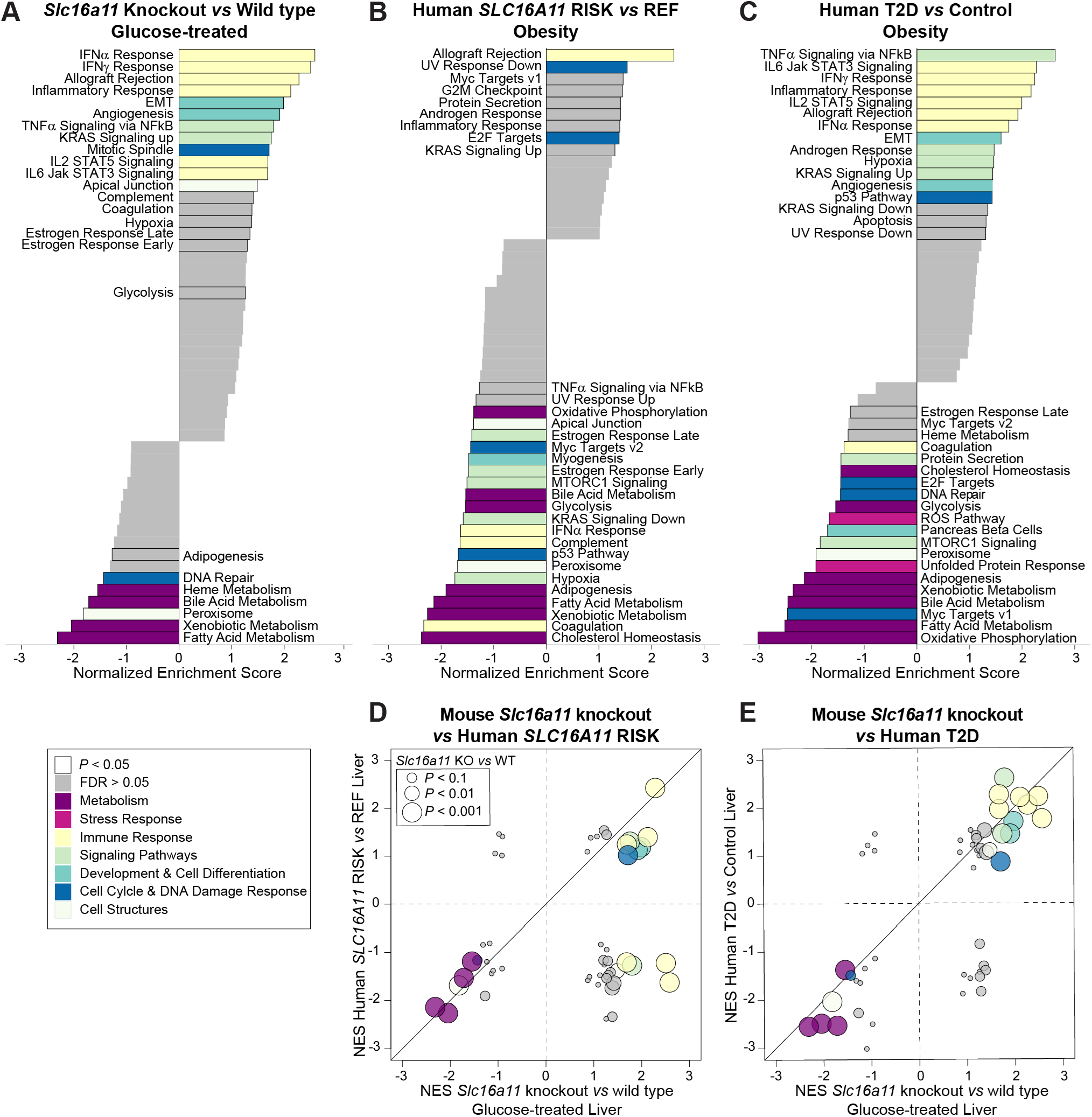
*Slc16a11* causes global changes in hepatic expression similar to what is observed in humans with T2D. (A-C) GSEA of liver transcriptomic profiles from (A) glucose-treated *Slc16a11*-knockout mice compared to wild-type littermates (n=5 each), (B) individuals without T2D who are homozygous T2D risk carriers (RISK) versus homozygous reference (REF) carriers of the *SLC16A11* haplotype (n=4 each), and (C) individuals with T2D versus controls (n=8 each). Each bar represents a different pathway from the Hallmark pathways collection (Liberzon et al., 2015). Pathways with *P* < 0.05 are labeled and outlined. Colors represent different categories of pathways. Non-significant pathways with (false discovery rate [FDR] > 0.05) are shown in gray. (D-E) Scatterplots comparing transcriptomic pathway changes observed in livers from glucose-treated *Slc16a11*-knockout mice with those observed in (D) human *SLC16A11* T2D risk haplotype carriers and (E) individuals with T2D. *P* values and FDR indicate results from comparison of glucose-treated *Slc16a11*-knockout *vs* wild-type mice. See also Figures S4, S5, and S6 and Tables S5 and S6.

The difference in expression profiles observed in fasting and glucose-treated mice is driven by the response of *Slc16a11*-deficient mice to glucose. In contrast to *Slc16a11*-deficient mice, wild-type mice show relatively few changes in pathway or zonal gene expression following glucose administration (*i.e.*, comparing glucose-treated wild-type mice to fasting wild-type mice (Figures S4C and S5A-E). Collectively, our data from fasting and glucose-treated mice suggest Mct11 deficiency leads to a level of chronic metabolic stress that predisposes the liver to an acute inflammatory response following nutrient stress, which is a process implicated in hepatic insulin resistance and T2D (Baker et al., 2011; Kim et al., 2015).

### Metabolic changes induced by Mct11 deficiency are observed in human *SLC16A11* T2D risk carriers and individuals with T2D

To demonstrate the relevance of changes observed in our *Slc16a11*-knockout mouse model to the human context, we also compared the transcriptomic profiles of liver biopsies from individuals who are homozygous carriers of the *SLC16A11* T2D risk and reference (non-risk) haplotypes as well as individuals with and without T2D. The individuals profiled were medication-free for 24 hours prior to sample collection, and show no differences in quantitative glycemic and lipid traits according to genotype or T2D status (Figures S6A-B). All individuals were obese (average BMI ∼40 kg/m^2^), and therefore exposed to chronic caloric excess and potentially nutrient stress.

To avoid the confounding effects of T2D, we assessed the impact of *SLC16A11* genotype in individuals without a T2D diagnosis. In comparison to individuals who carry the non-risk haplotype, liver biopsies of *SLC16A11* T2D-risk haplotype carriers show higher expression of genes involved in certain inflammatory and DNA damage response pathways, and lower expression of genes involved in metabolism, including cholesterol, bile acid, fatty acid, and xenobiotic metabolic processes and oxidative phosphorylation (Figure 6B and Table S6). Though there are differences in individual pathways, the overall impact of the T2D risk haplotype on expression in human liver is largely consistent with what we observed in our *Slc16a11* knockout mouse model.

Similarly, we found that individuals with T2D (including both carriers and non-carriers of the *SLC16A11* risk haplotype) have increased expression of genes involved in inflammatory response pathways, such as TNFα signaling via NFκB, IFNα and IFNγ response, signaling through IL6, IL2, and JAK-STAT, and EMT, and decreased levels of metabolic pathway gene expression, including oxidative phosphorylation, fatty acid metabolism, bile acid metabolism, and xenobiotic metabolism (Figure 6C and Table S6). These human physiological data are strikingly similar to what we observed in our glucose-treated *Slc16a11*-knockout mice (Figures 6D-E, comparing Figure 6A to 6B and 6C), and demonstrate that *Slc16a11* deficiency induces changes in hepatic metabolism consistent with what happens in individuals who carry the *SLC16A11* T2D risk haplotype and those with T2D.

To summarize, the *SLC16A11* T2D risk haplotype effect is due to disrupted function of MCT11—a crucial regulator of lactate levels and energy homeostasis in the liver—which leads to cellular and physiological changes in metabolism that are associated with insulin resistance and T2D in humans.

## DISCUSSION

Human genetic discoveries are a valuable and effective tool for target identification and therapeutic development, enabling a focus on biological processes validated to impact disease risk (Plenge et al., 2013). To date, numerous therapeutic approaches have been implemented based on genetic insights; these include treatments for monogenic diseases such as cystic fibrosis and Duchenne muscular dystrophy, and more complex disorders, including coronary artery disease and T2D (Cohen et al., 2006; Nauck, 2014; Robinson et al., 2015; Sabatine et al., 2015; Sheikh and Yokota, 2021; Wainwright et al., 2015). Though beneficial, the path from genetic association to clinical benefit is arduous, requiring in-depth investigation into the molecular, cellular, and physiological processes affected by the variants in order to identify and validate targets, model systems, and biomarkers for drug development. For most genetic associations, the causal gene has not yet been identified; and even among variants for which evidence supports a causal gene, the mechanisms leading to pathophysiology are rarely understood. Here, we follow up prior work that identified reduced MCT11 function in liver as the disease-relevant variant effect of the *SLC16A11* T2D risk haplotype (Rusu et al., 2017). We identify the low oxygen, pericentral environment as an essential context to understand variant-disrupted MCT11 function, and demonstrate that MCT11 deficiency alters glucose metabolism, lactate levels, and energy homeostasis, and leads to TAG accumulation. Our work suggests a mechanism whereby MCT11 deficiency increases T2D risk due to a shift in energy metabolism toward a state of nutrient excess, resulting in elevated metabolic stress and, ultimately, hepatic damage and dysfunction.

In addition to expanding our understanding of the *SLC16A11* disease mechanism, our discovery that *SLC16A11* expression has a pericentral zonation profile provides clarity on how MCT11 action is distinct from closely related family members. While further investigation is needed to confirm the role of MCT11 in lactate transport, our data suggest MCT11 may be the primary MCT responsible for lactate export from hepatocytes residing in the low oxygen pericentral region. This would be analogous to the role of *SLC16A3*/MCT4 in muscle, which is induced by hypoxia-inducible factor 1 (HIF1) and mediates lactate efflux under glycolytic, anaerobic conditions such as experienced during hypoxia and exercise (Bisetto et al., 2019; Contreras-Baeza et al., 2019; Ullah et al., 2006). Beyond the MCT family, our findings underscore the importance of delineating cell populations and contexts where genes are expressed and the value in modeling relevant cellular contexts when investigating gene function.

The *SLC16A11* pericentral zonation profile also offers insight into how the T2D risk haplotype decreases *SLC16A11* levels in liver, highlighting a disruption in oxygen tension-induced expression. However, the detailed mechanism through which the low oxygen pericentral environment regulates *SLC16A11* in hepatocytes and which T2D risk variant(s) disrupt this regulatory process are not yet known. The credible set for the T2D risk haplotype includes three noncoding variants located near the *SLC16A11* transcriptional start site (Rusu et al., 2017); however, none of these variants are predicted to overlap putative binding sites for hypoxia-inducible factors (HIFs) (unpublished analyses), suggesting *SLC16A11* could be regulated through HIF interactions with other variant-disrupted transcription factors or through a HIF-independent mechanism. Elucidating these regulatory processes is an important area for future study that would benefit translational efforts to counteract the haplotype effect.

Though our findings emphasize MCT11 action in pericentral hepatocytes, we do not know the relative contribution of *SLC16A11* disruption in pericentral *versus* periportal hepatocytes to T2D risk. We only detected lactate accumulation at low oxygen; however, *SLC16A11* deficiency also leads to TAG accumulation in PHH cultured at 21% oxygen (Rusu et al., 2017), an apparent discrepancy that could be explained by differences in metabolism kinetics or metabolic wiring in these environmental contexts. In mice, Mct11 deficiency has distinct effects on pericentral and periportal gene expression, with glucose exposure causing changes in zonal expression profiles similar to what occurs in models of liver fibrosis (Ghallab et al., 2019). Furthermore, despite its clear role in hepatocytes, the metabolic phenotypes caused by MCT11 deficiency may not be entirely due to its role in the liver. The discoveries that *SLC16A11* expression is regulated by oxygen and restricted within liver cell populations raise the possibility that *SLC16A11* might be similarly expressed in specific cell populations within other healthy tissues or in response to certain pathologies. For example, long-term high-fat diet feeding induces liver expression of *Slc16a11* in mice (Soltis et al., 2017; Zhang et al., 2019), an effect that might be due to obesity-induced changes in the liver oxygen gradient (Mantena et al., 2009).

Another aspect of the disease mechanism we do not yet fully understand is how the impact of MCT11 deficiency on lactate levels leads to metabolic phenotypes and increased T2D risk. Excess lactate serves as a nutrient signal that could promote hyperglycemia due to its inhibitory effect on glycolysis or through its role as a precursor for gluconeogenesis (Costa Leite et al., 2007; Leite et al., 2011). Likewise, TAG accumulation could result directly from glucose-derived carbons, from excess lactate being redirected through the glycerol-3-phosphate shuttle (which provides the lipid glycerol backbone), and/or from increased flux through the citrate shuttle driving fatty acid synthesis. The nutrient excess state generated by Mct11 deficiency emulates the effect of caloric excess, and may explain why *SLC16A11* risk haplotype carriers develop T2D at a lower BMI than individuals with T2D who are not *SLC16A11* risk carriers (SIGMA T2D Consortium et al., 2014). In addition, metabolic and nutrient stress are signals that lead to hepatic damage, inflammation, and insulin resistance (Chen et al., 2015; Del Campo et al., 2018; Tanti et al., 2012), consistent with evidence that human *SLC16A11* carriers have signs of hepatic damage and decreased insulin action (Almeda-Valdes et al., 2019).

Though *Slc16a11* deficiency does not cause robust changes in glucose homeostasis in mice (Hoch et al., 2019; Zhao et al., 2019a; Zhao et al., 2019b), our current and previous studies provide clear evidence that human *SLC16A11* T2D risk variants reduce MCT11 function, and that MCT11 deficiency causes risk haplotype- and T2D-relevant phenotypes—supporting prior hypotheses that therapeutic approaches that increase *SLC16A11* expression and MCT11 function may be beneficial for individuals with T2D (Rusu et al., 2017). This work also provides key pieces of information critical for advancing therapeutic development, including identifying lactate as a potential circulating biomarker of effect that could be used to assess activity *in vivo*. Evaluation of lactate levels in glycemic studies of human *SLC16A11* carriers, as well as in T2D risk variant cell and animal models, is a crucial next step in validating this effect.

While our data suggest a favorable metabolic effect from increasing MCT11 function, efforts to do so will require precision and specificity. Endogenous *SLC16A11* has a restricted expression profile, which raises the possibility that therapeutic approaches may need to target the pericentral region to prevent metabolic effects that can arise from misexpression. Furthermore, while higher *SLC16A11*/MCT11 expression is associated with improved survival in liver, renal, and pancreatic cancers (The Human Protein Atlas SLC16A11), increased expression of both *SLC16A1*/MCT1 and *SLC16A3*/MCT4 are negative prognostic indicators for cancer survival (The Human Protein Atlas SLC16A1; The Human Protein Atlas SLC16A3; Uhlen et al., 2017), and inhibition of lactate transport mediated by MCT1 and MCT4 is currently being investigated as a treatment approach for cancer (Doherty and Cleveland, 2013). This suggests that increasing MCT11 activity is not expected to increase cancer risk but highlights the need for specificity, which can be accomplished through targeted gene therapy or other precise approaches that correct the haplotype effect. In conclusion, this study demonstrates how in-depth functional interrogation of genetic risk variants can reveal mechanisms contributing to disease pathophysiology, and support development of therapeutics with potential to benefit individuals with T2D.

## ACKNOWLEDGEMENTS

The authors thank Melina Claussnitzer and Bridget Wagner for many helpful discussions and critical reading of the manuscript. We also acknowledge the many discussions with the Florez and Wagner laboratories and the Diabetes Research Group at the Broad Institute. Several depictions were created using images available from BioRender.com. This work was conducted as part of the Slim Initiative for Genomic Medicine, a project funded by the Carlos Slim Foundation in Mexico, and with support from the Next Generation Fund at the Broad Institute of MIT and Harvard to S.B.R.J.

## AUTHOR CONTRIBUTIONS

A.A., L.Z., V.R., J.C.F., and S.B.R.J. conceived and planned the study. A.A., L.Z., V.R., and S.B.R.J. designed experiments. A.A. performed experiments in primary human hepatocytes. V.R. designed the CRISPR targeting approach and sequenced the targeted region to generate the *Slc16a11^del19^* knockout mouse model. P.G., K.G.-R., and L.Z. carried out animal husbandry, genotyping, and mouse studies. J.C. conducted the TSH and T4 assays, which A.N.H. supervised. P.G., L.Z., A.A., and S.B.R.J. analyzed data from mouse studies. A.A and V.R. performed RNA sequencing. A.A., V.R., and S.B.R.J analyzed RNA-sequencing data. A.D., K.P., K.B., C.D., S.J., and J.K. processed metabolite profiling samples by mass spectrometry, which C.C. supervised. S.B.R.J analyzed the resulting metabolite profiling datasets. A.H provided technical support. E.H. assisted with writing. F.C-C., F.B.-O., C.Z., and L.O. provided tissue samples. E.S.L., J.C.F., and S.B.R.J. oversaw the study. A.A., J.C.F., and S.B.R.J. wrote the manuscript.

## RESOURCE AVAILABILITY

### Lead contact

Further information and requests for resources and reagents should be directed to and will be fulfilled by Suzanne Jacobs.

### Materials and data availability

Certain materials are shared with academic and non-profit research organizations for research and educational purposes only under an MTA to be discussed in good faith with the recipient. The RNA sequencing data generated in this study will be deposited in the NCBI GEO database, and will be publicly available upon publication of this manuscript. This paper does not report original code.

## METHODS

### Human Subjects

Liver tissue collection and study participants have been described previously (Rusu et al., 2017). All contributing studies were approved by their respective local ethics committees.

### Primary Human Hepatocytes

Primary human hepatocytes (PHH) were purchased from BioIVT. Genotyping of two SNPs on the *SLC16A11* T2D risk haplotype (rs13342232 and rs75493593) was carried out by BioIVT and used to determine homozygosity at the *SLC16A11* locus. Donor lots used in gene expression studies include YNZ, BHL, VHB, and RSF (homozygous for the non-risk, reference haplotype) and YNS and VDT (homozygous for the T2D risk haplotype; carriers of the alternative allele at both SNPs). Metabolite profiling experiments used donor lot YNZ. Cells were thawed and immediately resuspended in CP media (BioIVT) supplemented with torpedo antibiotic (BioIVT). Cell count and viability were assessed by trypan blue exclusion test prior to plating.

Hepatocytes were plated onto collagen-coated 24 well plates (Corning, 354408) or 96 well plates (Corning, 354407) at a density of 350,000 or 50,000 cells per well, respectively, in CP media supplemented with torpedo antibiotic (BioIVT). After 4 h media was replaced with fresh CP or HI media, depending on the experiment. Cells were then maintained in 21% oxygen/5% CO_2_ or moved to a Thermo Heracell 240i Tri-Gas incubator and maintained at 4% oxygen/5% CO_2_. After 24 h, media was replaced with fresh CP or HI media. Hepatocytes were grown for an additional 24 h prior to further processing.

### *Slc16a11*-Knockout Mice

All animal work was approved by the Institutional Animal Care and Use Committee (IACUC) at the Broad Institute and carried out in accordance with local, state, and federal regulations. We targeted the *Slc16a11* allele for inactivation using CRISPR-Cas9 technology. An *in vitro* transcribed sgRNA targeting *Slc16a11* (5’-AGTCCTAACCTCGCTTGGCT-3’) and hspCas9 mRNA (CAS500A-1, System Biosciences) were microinjected into C57BL/6J mouse zygotes by the Genome Modification Facility at Harvard University. We used next-generation sequencing to compare the targeted region of *Slc16a11* CRISPR-Cas9 mice to the wild-type sequence and identified two chimeric mice with the same 19 bp deletion (resulting in a frameshift leading to early termination and a truncated Mct11 protein). These founders were bred to C57BL/6J mice to create germline *Slc16a11^del19^* lines, and then heterozygous *Slc16a11^del19^* progeny from each line were intercrossed to remove potential non-linked, off-target effects. All animals were maintained on a C57BL/6J background.

### Topology

The membrane topology of mouse Mct11 (Uniprot ID Q5NC32) and the protein encoded by the *Slc16a11^del19^* frameshift were determined using the Constrained Consensus TOPology prediction server (http://cctop.enzim.ttk.mta.hu/?_=) and visualized with TexTopo (Beitz, 2000).

### Animal Care and Husbandry

To generate wild-type and *Slc16a11*-knockout littermates for experiments, *Slc16a11^del19^* heterozygous mice were intercrossed. Littermates were housed together in a pathogen free facility with a 12:12 light:dark cycle. Animals used for experiments were kept on wood bedding beginning at four weeks of age, and fed irradiated normal chow diet (PicoLab Rodent Diet 20, 5053). Body composition was measured using quantitative nuclear magnetic resonance spectroscopy (EchoMRI Analyzer, Echo Medical Systems) in live, conscious animals.

### Mouse Genotyping

Mice were genotyped using tissue collected from an ear punch. Genomic DNA was extracted by incubating the tissue with proteinase K buffer [1% 0.5 M EDTA, 1% 20% SDS, 4% 5M NaCl, 10% 1M Tris HCl pH 7.6, distilled H2O, 1% proteinase K (20mg/mL)] in a Thermomixer set at 55C and 1200 rpm for 30 min. Insoluble material was pelleted and DNA was precipitated from supernatant using equal volume isopropanol. DNA was solubilized in water. Genotyping was performed by ddPCR using the ddPCR Supermix for Probes (No dUTP) kit (Bio-Rad) and the following primer and probe sets: wild-type allele (Fwd primer: 5’-ACTCTTCTTACCAGCGAG-3’; Rev primer: 5’-GGGGGAGTCCTAACCT-3’; HEX-labeled probe 5’-CGAGAAGACCAAGCCAAG-3’) and *Slc16a11^del19^* allele (Fwd primer: 5’-ACTCTTCTTACCAGCGAG-3’; Rev primer: 5-GTGATGGTTGGGGGAG-3’; FAM-labeled probe 5’-GGCGAAAGCGAGGTTA-3’). Droplets were generated and analyzed using a QX200 Droplet Generator and Reader system (Bio-Rad). Data was extracted using QuantaSoft (Bio-Rad) and analyzed using Microsoft Excel. Genotype was determined by the percentages of wild-type and knockout alleles.

### Gene Expression Analyses by Droplet Digital PCR (ddPCR)

Total RNA was extracted using the RNeasy Mini Kit (Qiagen) or RNAClean XP beads (Life Sciences) and DNase treated prior to measuring expression. Target quantification was performed using the One-Step RT-ddPCR Advanced Kit for Probes (BioRad) and FAM- or HEX-labeled, TaqMan Real-Time PCR assays (IDT) as detailed below. Droplets were generated and analyzed using a QX200 Droplet Generator and Reader system (BioRad). Data was extracted using QuantaSoft (BioRad) and analyzed using Microsoft Excel. Expression counts in mouse tissues were adjusted using normalization factors calculated from *Tbp*, and shown as either normalized counts or levels relative to wild type. Expression levels in PHH were normalized to *TBP* and are shown as levels relative to the indicated control. Human gene expression studies included 3-8 biological replicates from each donor per experiment. *P* values were computed using a Student’s t test between two sample types.

| Target Gene | Organism | Assay ID | Fluorophore | Source |
| --- | --- | --- | --- | --- |
| *SLC16A11* | HOMO_SAPIENS | Hs.PT.58.4710484.g | FAM | IDT |
| *SLC16A13* | HOMO_SAPIENS | Hs.PT.58.22429467 | FAM | IDT |
| *TBP* | HOMO_SAPIENS | Hs.PT.39a.22214825 | HEX | IDT |
| *Slc16a1* | MUS_MUSCULUS | Mm.PT.58.7462799 | FAM | IDT |
| *Slc16a2* | MUS_MUSCULUS | Mm.PT.58.42920302 | FAM | IDT |
| *Slc16a3* | MUS_MUSCULUS | Mm.PT.58.43168758.g | FAM | IDT |
| *Slc16a4* | MUS_MUSCULUS | Mm.PT.58.43799834 | FAM | IDT |
| *Slc16a5* | MUS_MUSCULUS | Mm.PT.58.16984913 | FAM | IDT |
| *Slc16a6* | MUS_MUSCULUS | Mm.PT.58.31075711 | FAM | IDT |
| *Slc16a7* | MUS_MUSCULUS | Mm.PT.58.14123799 | FAM | IDT |
| *Slc16a8* | MUS_MUSCULUS | Mm.PT.58.42641245 | FAM | IDT |
| *Slc16a9* | MUS_MUSCULUS | Mm.PT.56a.13207630 | FAM | IDT |
| *Slc16a10* | MUS_MUSCULUS | Mm.PT.58.31920361 | FAM | IDT |
| *Slc16a11* | MUS_MUSCULUS | Mm.PT.58.7890237 | FAM | IDT |
| *Slc16a12* | MUS_MUSCULUS | Mm.PT.58.32850803 | FAM | IDT |
| *Slc16a13* | MUS_MUSCULUS | Mm01250819_m1 | FAM | Thermo |
| *Slc16a14* | MUS_MUSCULUS | Mm.PT.58.13602391 | FAM | IDT |
| *Bcl6b* | MUS_MUSCULUS | Mm00455912_g1 | FAM | Thermo |
| *Tbp* | MUS_MUSCULUS | Mm.PT.39a.22214839 | HEX | IDT |

### Glucose Tolerance Tests

The evening prior to each intraperitoneal glucose tolerance test (IPGTT), mice were housed in a fresh cage without food but with water. Following a ∼16 h overnight fast, the mice were weighed and fasting glucose (time 0 min) was measured in blood collected from the tail vein using an AlphaTrak 2 glucometer (animal code 14). Glucose (1 or 1.5 g/kg in PBS) was administered by intraperitoneal injection and blood glucose levels were measured at 15, 30, 60, and 120 min. At the 0 and 30 min time points of some experiments, the tail was briefly warmed with a heat lamp to enable additional blood collection for insulin measurements. Following completion of the IPGTT, animals were returned to their home cage with normal enrichment and food. Some experiments were terminated early for tissue collection.

### Blood and Tissue Collection

For terminal blood and tissue collection, mice were anesthetized by isoflurane inhalation. Blood was collected by retro-orbital eye bleeding into an EDTA KE/1.3 tube for plasma. Tissues were harvested quickly and flash frozen in liquid nitrogen for storage at −80C or fixed for histology. For gene expression and metabolite analyses, frozen tissue was crushed into a powder using a Covaris CP02 cryoPREP tissue pulverizer. Powdered tissue was stored at −80C prior to subsequent gene expression and metabolite analyses. For histology, tissue samples were fixed in 4% paraformaldehyde, washed with PBS, and stored in 70% ethanol until paraffin embedding.

### Measurements of Blood Samples

Insulin was measured in plasma samples (10 µL) collected from mice at the 0 (fasting) and 30 min time points of an IPGTT using a rat/mouse insulin ELISA kit following manufacturer’s instructions (Ultra Sensitive Insulin ELISA Kit; Crystal Chem Inc, Catalog #90080). Samples with values below the detectable limit of the assay were excluded from downstream analyses. Liver function was assessed by measuring serum alanine aminotransferase (ALT) activity using an ALT fluorometric Assay Kit (BioVision, Cat# K752-100). Total T4 (TT4) levels were measured using AccuDiag^™^ T4 ELISA Kit Cat# 3149–18 (Diagnostic Automation /Cortez Diagnostics, Inc. Calabasas, CA) in 25 µL of serum. Circulating thyroid-stimulating hormone (TSH) was measured in 10 μL serum via Milliplex MAP Mouse Pituitary Magnetic Bead Panel Mouse (EMD Millipore, Billerica, MA) on a Luminex 200 (Luminex Corporation, Austin, TX).

### Analyses of Mouse Physiological Data

Physiological data from mouse studies, including measurements of body weight, body composition, glucose, insulin, ALT, T4, and TSH, underwent quality control (QC) and analysis as follows. For all measurements, individual data points were removed if they were more than 2 standard deviations from the mean value for that measurement within each genotype. Glucose area-under-curve (AUC) calculations only include mice with data at all IPGTT time points following outlier removal. Insulinogenic index (IGI) calculations only include mice with glucose and insulin data at both 0 and 30 min timepoints following outlier removal. *P* values were computed using a Wilcoxon rank sum test between wild-type and *Slc16a11*-knockout mice.

### Histological Analyses

Hematoxylin and Eosin (H&E) and picrosirius red staining on PFA-fixed, paraffin-embedded tissues was performed by the Histology Core at Beth Israel Deaconess Medical Center. Images were captured using a Zeiss Observer Z1 microscope using a 10x eyepiece with 20x-63x objectives, and ZEN Pro v2.0.0 software.

### Preparation of Samples for Metabolite Profiling

In primary human hepatocytes (donor lot YNZ), knockdown of *SLC16A11* expression was achieved using the Accell Human SLC16A11 siRNA SMARTpool (E-007404-00-0050, GE Dharmacon). The Accell Non-targeting Pool (D-001910-10-50, GE Dharmacon) was used as a control. siRNAs were reconstituted in RNase-free 1x siRNA Buffer at 100 μM. Hepatocytes were plated into collagen-coated 24 well plates (Corning, 354408) at a density of 350,000 cells per well. After 4 hours, cells were washed once with HI media (BioIVT) and siRNAs in HI media were added at a final concentration of 1 μM. Cells were then maintained at 21% oxygen or moved to a Thermo Heracell 240i Tri-Gas incubator at 4% oxygen. After 24 h, media was replaced with fresh CP media and hepatocytes were grown for another 24 h, after which the baseline (t=0) samples were collected. For time points of 15 min, 2 h, and 6 h stimulation, media was replaced with fresh CP media containing 1g/L glucose and samples were collected for metabolite profiling after the indicated time. At each time point and at each oxygen tension, three biological replicates were for collected for siRNAs targeting *SLC16A11* and for negative controls. Due to the media change, baseline (t=0) differences in extracellular metabolites reflect changes accumulated during the 24 h period prior to sample collection.

Since extraction of lipid and polar metabolites from cells requires different solutions, we generated duplicate sets of hepatocyte samples that were collected in parallel. For sample collection, primary human hepatocytes were removed from the incubator and immediately placed on ice. Media (500 µL) was collected from the hepatocytes to assess extracellular metabolite changes. Hepatocytes were then washed with 500 µL ice cold PBS. For analysis of lipids, metabolites were extracted by scraping hepatocytes in 320 µL of isopropanol (HPLC Grade; Honeywell) containing 1,2-didodecanoyl-sn-glycero-3-phosphocholine (Avanti Polar Lipids; Alabaster, AL). For analysis of polar metabolites and free fatty acids, metabolites were extracted by scraping hepatocytes in 320 µL of 80% methanol (VWR) containing 0.05 ng/µL inosine-15N4, 0.05 ng/µL thymine-d4, and 0.1 ng/µL glycocholate-d4 as internal standards (Cambridge Isotope Laboratories). Media was spun at 600 g at 4C for 5 min to remove any debris.

For mouse samples, metabolite profiling was performed on liver and plasma collected from overnight-fasted mice following administration of PBS (n=6 wild-type mice and n=4 *Slc16a11* knockout) or 1.5 g/kg glucose (n=5 each wild type and *Slc16a11* knockout) for 30 minutes. Plasma was collected as described above. Powdered liver tissue from *Slc16a11* knockout and wild-type mice was homogenized in a weight-proportional volume 1:4 with water. Tungsten beads (2x 3mm, Qiagen) were added to each sample and homogenized in a Tissuelyser II (Qiagen) set at 20Hz for 4 minutes. Homogenates were then aliquoted in 10 and 30 µL aliquots for each method.

Metabolites from PHH media, mouse tissue homogenates, and mouse plasma were precipitated for lipid (C8-pos) analyses by taking 10 µL sample and adding 190 µL isopropanol. Metabolites were precipitated for free fatty acid (C18-neg) analyses using 30 µL sample + 90 µL extraction solution (0.05 ng/µL 15R-15-methyl ProstaglandinA2, 0.05 ng/µL 15R-15-methyl ProstaglandinF2α, 0.05 ng/µL 15R-15S-15-methyl ProstaglandinD2, 0.05 ng/µL 15S-15-methyl ProstaglandinE1, 0.05 ng/µL 15S-15-methyl ProstaglandinE2 in Methanol). Polar metabolites were precipitated for HLIC-neg with 30 µL sample + 120 µL extraction solution (0.05 ng/µL Inosine-15N4, 0.05 ng/µL Thymine-d4, 0.1 ng/µL Glycocholate-d4 in 80% Methanol). Finally, polar metabolites were precipitated for HILIC-pos using 10 µL sample + 90 µL extraction solution consisting of Acetonitrile:Methanol:Formic acid (75:25,0.2 v:v:v) with a nominal concentration of 0.2 µg/mL valine-d8 (Sigma) and phenylalaine-d8 (Cambridge Isotopes Laboratories).

### Metabolite Profiling

Analyses of lipids were conducted using an LC-MS system comprised of a Shimadzu Nexera X2 U-HPLC (Shimadzu Corp.; Marlborough, MA) coupled to an Exactive Plus orbitrap mass spectrometer (Thermo Fisher Scientific; Waltham, MA). Lipid extracts were injected onto an ACQUITY BEH C8 column (100 x 2.1 mm, 1.7 µm; Waters, Milford, MA). The column was eluted isocratically with 80% mobile phase A (95:5:0.1 vol/vol/vol 10 mM ammonium acetate/methanol/formic acid) for 1 minute followed by a linear gradient to 80% mobile-phase B (99.9:0.1 vol/vol methanol/formic acid) over 2 minutes, a linear gradient to 100% mobile phase B over 7 minutes, then 3 minutes at 100% mobile-phase B. MS data were acquired using electrospray ionization in the positive ion mode over 220–1100 *m/z* and at 70,000 resolution. Other MS settings were: sheath gas 50, in source CID 5 eV, sweep gas 5, spray voltage 3 kV, capillary temperature 300°C, S-lens RF 60, heater temperature 300°C, microscans 1, automatic gain control target 1e6, and maximum ion time 100 ms. Raw data data were processed using TraceFinder 3.3 (Thermo Fisher Scientific; Waltham, MA) and Progenesis QI (Nonlinear Dynamics; Newcastle upon Tyne, UK) software for detection and integration of LC-MS peaks. Lipid identities were determined based on comparison to reference standards and reference plasma extracts and are denoted by total number of carbons in the lipid acyl chain(s) and total number of double bonds in the lipid acyl chain(s). Negative ion mode analyses of free fatty acids and bile acids (C18-neg) were conducted using an LC-MS system composed of a Shimadzu Nexera X2 U-HPLC (Shimadzu Corp) coupled to a Q Exactive hybrid quadrupole orbitrap mass spectrometer (Thermo Fisher Scientific). The samples were injected onto a 150×2.1 mm, 1.7 um ACQUITY BEH C18 column (Waters; Milford, MA). The column was eluted isocratically at a flow rate of 450 µl/min with 80% mobile phase A (0.01% formic acid in water) for 3 minutes followed by a linear gradient to 100% mobile phase B (acetonitrile with 0.01% acetic acid) over 12 minutes and held for 3 minutes. Column re-equilibration at initial conditions were performed for 8 minutes. MS analyses were performed in the negative ion mode using electrospray ionization, full scan MS acquisition over 70 to 850 m/z, and a resolution setting of 70000. Metabolite identities were confirmed using authentic reference standards. Other MS settings were as follows: sheath gas 45, sweep gas 5, spray voltage −3.5 kV, capillary temperature 320°C, S-lens RF 60, heater temperature 300°C, microscans 1, automatic gain control target 1e6, and maximum ion time 250 ms. Samples for negative and positive ion mode analyses of polar metabolites were achieved using the HILIC (hydrophilic interaction chromatography) method under basic conditions as described previously (SIGMA T2D Consortium et al., 2014). Metabolites of unknown identity were excluded from analyses.

### Analysis of Metabolite Profiling Data from Primary Human Hepatocytes

For each PHH time point, metabolite profiling data across both oxygen tensions were first total signal normalized within each of the 4 LC-MS methods. Data were log2 transformed and then filtered to remove duplicate metabolites. The ratio of glutamine to glutamate and total levels of select lipid subclasses were calculated, and then individual metabolite data were removed if they were more than 2 standard deviations from the mean value within a sample type (where sample types are *SLC16A11* siRNA or control siRNA treatment within each time point and oxygen tension). Metabolites with fewer than two replicates in each sample type were removed. For comparisons of *SLC16A11* knockdown versus control, metabolite abundances were normalized to the median value for that metabolite within the control siRNA replicates. Fold changes for each metabolite were calculated using the median normalized value across *SLC16A11* siRNA samples and the median normalized value across control siRNA samples. To compare metabolite abundance at 4% and 21% oxygen, metabolite abundances in control siRNA samples at each oxygen tension were normalized to the median value for that metabolite within the 21% oxygen control siRNA samples. Fold changes for each metabolite were calculated using the median normalized value across control siRNA samples at 4% oxygen and the median normalized value across control siRNA samples at 21% oxygen. *P* values were computed using a Student’s t test between two sample types.

### Analysis of Metabolite Profiling Data from *Slc16a11*-knockout and wild-type mice

For each tissue, metabolite-profiling data across treatment conditions (i.e., fasting and glucose treated) were first total signal normalized within each of the 4 LC-MS methods. Data were log2 transformed and then filtered to remove duplicate metabolites. The ratio of glutamine to glutamate and total levels of select lipid subclasses were calculated, and then individual metabolite data were removed if they were more than 2 standard deviations from the mean value within a sample type (where sample types are *Slc16a11* knockout or wild type within each tissue and treatment condition). Metabolites with fewer than two replicates in each sample type were removed. For comparisons of *Slc16a11* knockout versus wild type, metabolite abundances were normalized to the median value for that metabolite within the wild type replicates. Fold changes for each metabolite were calculated using the median normalized value across *Slc16a11* knockout samples and the median normalized value across wild type samples. To compare metabolite abundance in fasting and glucose-treated mice, metabolite abundances were normalized to the median value for that metabolite within the fasting samples. Fold changes for each metabolite were calculated using the median normalized value across glucose-treated samples and the median normalized value across wild-type samples. *P* values were computed using a Student’s t test between two sample types.

### Metabolite Pathway Analysis

To identify metabolite changes at the pathway level, we applied a strategy that is commonly used for analysis of gene expression data: gene-set enrichment analysis (GSEA) (Mootha et al., 2003; Subramanian et al., 2005). Pathway enrichment was computed using the GSEA v3.0 PreRanked tool, as implemented at http://software.broadinstitute.org/gsea/index.jsp, using an unweighted enrichment score and 1,000 permutations. The log2-transformed fold changes were used as input, along with curated sets of 47 KEGG pathways from the human reference set and 26 additional classes of metabolites covering bile acid, fatty acid, and lipid sub-types and acylcarnitines. Bile acid and fatty acid subclasses were identified using annotations from the Human Metabolome Database (HMDB). Only metabolite pathways and classes with at least 5 members measured in our dataset were considered.

### RNA-sequencing and Data Analysis

RNA-sequencing was performed on liver samples from overnight-fasted mice following administration of PBS (n=6 wild-type mice and n=4 *Slc16a11* knockout) or 1.5 g/kg glucose (n=5 each wild type and *Slc16a11* knockout) for 30 minutes. Total RNA was extracted from mouse liver samples using RNeasy Mini Kit (Qiagen), DNase treated and 1 μg total RNA was processed for mRNA sequencing. The sequencing library construction was performed with TruSeq Stranded mRNA kit (Illumina) according to the manufacturer’s protocol. Library quality was assessed with BioAnalyzer (Agilent) and concentration was determined by Qubit fluorometric quantification (Thermo Fischer Scientific). Libraries were sequenced on an Illumina NextSeq 500/550 High Output Kit v2.5 (75 Cycles) paired end.

Human liver samples were collected from subjects undergoing bariatric surgery for severe obesity (BMI greater than 40 kg/m2, or greater than 35 kg/m2 with comorbid entities) or elective surgery in non-obese patients. All individuals were Mexican Mestizos older than 18 years, carefully selected from the Integral Clinic of Surgery for Obesity and Metabolic Diseases or General Surgery Department at the Tláhuac Hospital in Mexico City. Liver biopsy was obtained at the distal end of the left hepatic lobe, just above the spleen. Samples were frozen immediately after removal. The protocols for collecting the liver samples was approved by the respective local research and ethics committees and all patients signed an informed consent. The Broad Genomics Platform extracted RNA from frozen tissue samples using the miRNeasy Mini Kit (Qiagen).

RNA-sequencing was performed on 8 liver samples from individuals with T2D and 8 individuals without T2D. Both the T2D and non-T2D groups include 4 individuals homozygous for the *SLC16A11* reference haplotype and 4 individuals homozygous for the T2D risk haplotype. All individuals are female and there were no significant differences in age, BMI, fasting glucose, total cholesterol, triglycerides, HDL cholesterol, insulin, Hb1Ac, systolic BP, or diastolic BP at the time of sample collection (Figure S6), as evaluated using analysis of variance followed by Tukey’s HSD post-hoc test. Total RNA (250 ng) was depleted for ribosomal RNA using NEBNext® rRNA DepletionKit (New England BioLabs) according to the manufacturer’s protocol. First strand synthesis was completed with Maxima Reverse Transcriptase using random hexamers (Thermo Fisher Scientific). Second strand synthesis was done with NEBNext® mRNA Second Strand Synthesis Module (New England BioLabs). Illumina library construction was performed with Nextera® XT DNA Library Preparation Kit (Illumina) according to the manufacturer’s protocol with the following modification: 0.4 ng of dsDNA was tagmented at 55°C for 10 min. Library quality was assessed with BioAnalyzer (Agilent) and concentration was determined by Qubit fluorometric quantification (Thermo Fischer Scientific). Libraries were sequenced on an Illumina NextSeq500, 75 bp paired end.

We assessed read quality in both mouse and human RNA-sequencing datasets using Fastqc (http://www.bioinformatics.babraham.ac.uk/projects/fastqc/). STAR was used to map reads to an index generated from GRCm38 and Gencode vM22.primary_assembly (for mouse data) and GRCh38 and Gencode v32.primary_assembly (for human data) and to summarize the reads across genes (Dobin et al., 2013). Finally, DESeq2 was used to perform differential expression analysis (Love et al., 2014).

### RNA-sequencing Pathway Analyses

For both mouse and human RNA-sequencing datasets, we used GSEA to identify transcriptional changes in 50 pathway gene sets that comprise the Hallmark pathway collection in MsigDB (Liberzon et al., 2015; Mootha et al., 2003; Subramanian et al., 2005). Pathway enrichment was computed based on log2-transformed fold changes using the GSEA v3.0 PreRanked tool, as implemented at http://software.broadinstitute.org/gsea/index.jsp, using a weighted enrichment score and 1,000 permutations.

### Analyses of Zonal Gene Expression in RNA-sequencing Data

We used GSEA to identify transcriptional changes in genes with pericentral and periportal zonation profiles. Four gene sets were constructed based on zonation profiles from single-cell RNA-sequencing analyses of mouse liver (Halpern et al., 2017). The pericentral and periportal gene sets includes the top 100 genes (sorted based on *P* value) with either a pericentral or periportal expression profile, respectively, from Supplementary Table 3 from Halpern et al., 2017. Two gene sets consisting of 100 genes each were constructed from genes without zonal expression profiles (bottom of the gene list when sorted by *P* value) and used as negative controls. Pathway enrichment in mouse liver RNA-sequencing data was computed based on log2-transformed fold changes using the GSEA v3.0 PreRanked tool using a weighted enrichment score and 1,000 permutations.

### *SLC16A11* Zonation Profiles in Mouse and Human Liver

Zonation and cell type expression profiles for *Slc16a11*, *Slc16a13*, and *Slc16a1* in mouse liver were obtained from published single-cell RNA-sequencing analyses [Supplementary Table 3 (Halpern et al., 2017) and Supplementary Data 2 (Halpern et al., 2018)]. Human zonation profiles for *SLC16A11*, *SLC16A13*, and *SLC16A1* were generated from published laser-capture microdissection (LCM) RNA-sequencing analyses of human liver (GEO dataset GSE83990) (McEnerney et al., 2017). The ratio of pericentral to periportal expression in the human liver LCM RNA-sequencing data was calculated using the mean normalized counts for each zone. *P* values were calculated using a Student’s t test.

## SUPPLEMENTAL FIGURES and FIGURE LEGENDS

**Figure S1. Related to Figure 2.**
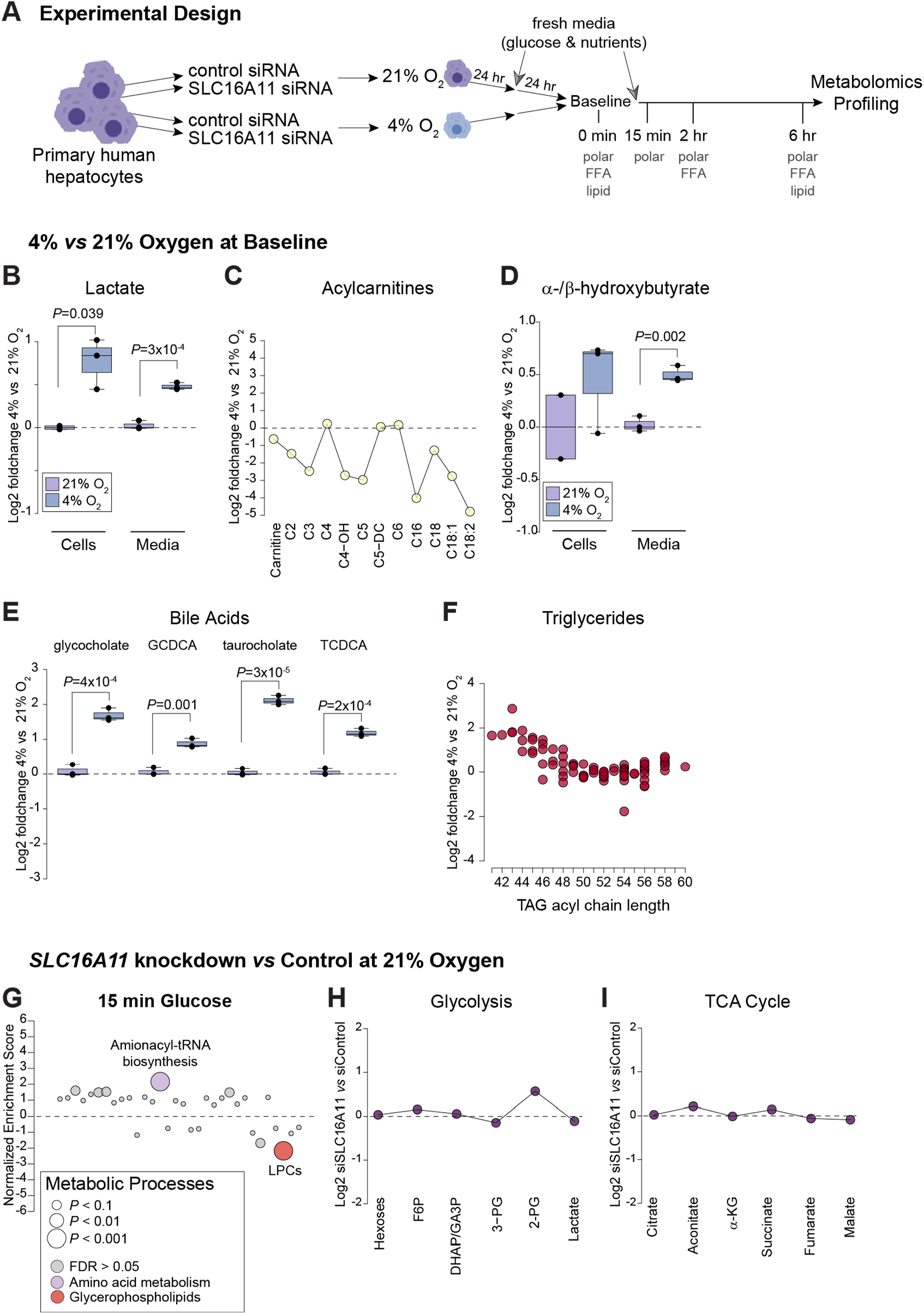
Metabolic changes induced by low oxygen and MCT11 deficiency in primary human hepatocytes. (A) Depiction of experimental design and timeline to assess the impact of *SLC16A11* knockdown on glucose metabolism of primary human hepatocytes (PHH) cultured under standard 21% oxygen tension and physiologic 4% pericentral oxygen tension. Categories of metabolites evaluated at each time point are indicated. (B-F) Effect of 4% oxygen tension on baseline levels of (B) lactate, (C) acylcarnitines, (D) ketone bodies, (E) bile acids, and (F) TAGs. Individual TAGs are grouped by acyl chain length. Plots depict log2 foldchange relative to control PHH cultured at 21% oxygen. (G-I) Intracellular metabolic pathway changes in *SLC16A11* knockdown PHH cultured under 21% oxygen tension and stimulated with fresh glucose-containing media for 15 minutes. (G) Pathway enrichment analysis. Each dot represents a different metabolic pathway or metabolite class. *P* values are indicated by dot size. Significantly altered pathways and metabolite classes (false discovery rate [FDR] < 0.05) are labeled and colored according to KEGG metabolic process or metabolite class, with non-significant pathways shown in gray. LPCs, lysophosphatidylcholines. (H-I) Line plots showing log2 fold changes in individual metabolites in (H) glycolysis and (I) the TCA cycle.

**Figure S2. Related to Figure 3.**
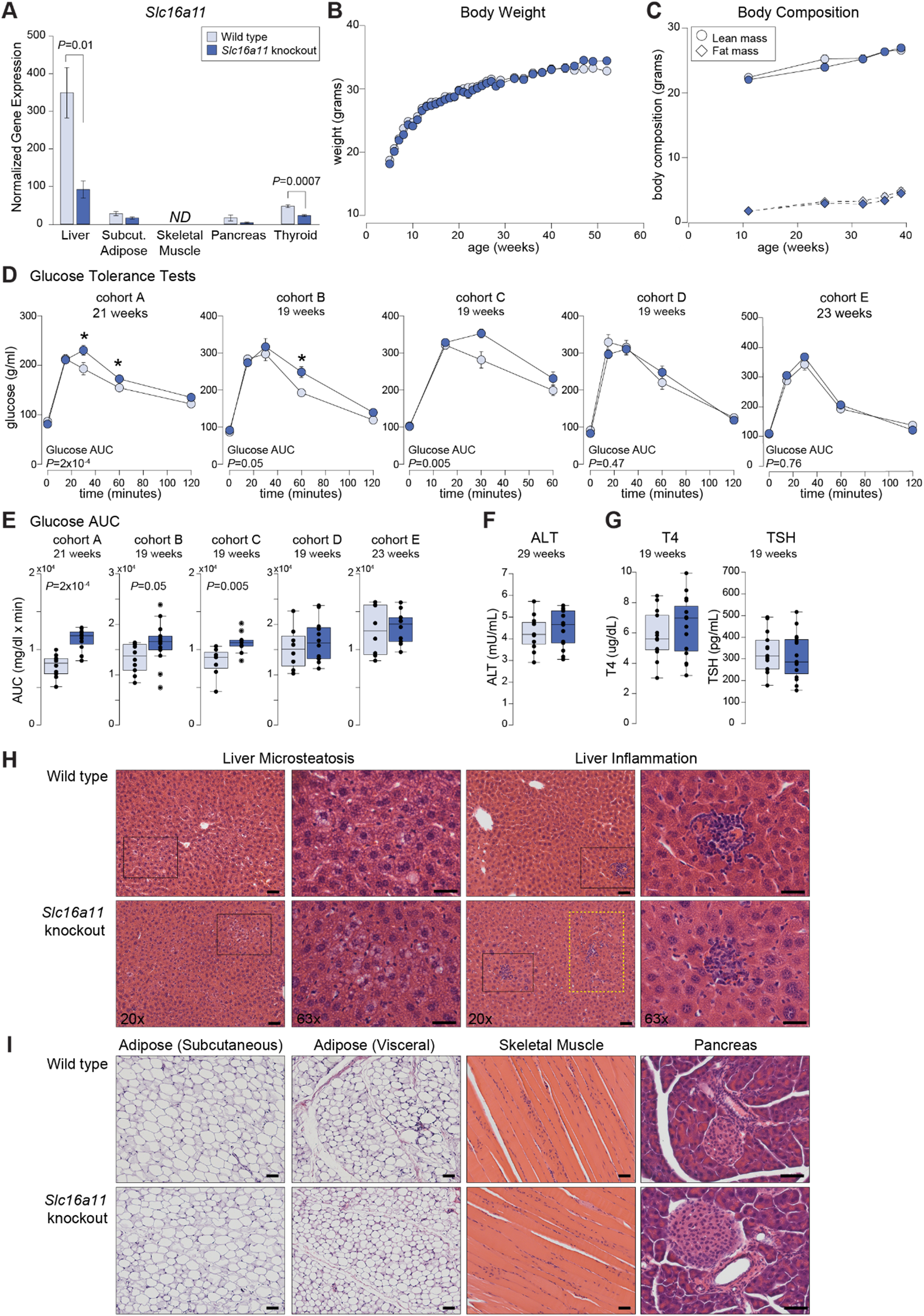
Physiological effects of Mct11 deficiency. (A) *Slc16a11* expression in tissues from adult wild-type (n=5) and *Slc16a11* homozygous knockout (n=4-5) mice. Bar plots depict relative gene expression ± SEM using *TBP* for normalization. *ND*=not detected. (B) Body weight measured over time. n=11-13 wild type and n=14-16 *Slc16a11* homozygous knockout mice. (C) Body composition (lean mass and fat mass) measured over time. n=12-13 wild type and n=13-14 *Slc16a11*-knockout mice per week. (D-E) Example IPGTTs and glucose area-under-curve boxplots from five independent cohorts of wild-type and *Slc16a11*-knockout mice between 19-23 weeks of age. (F) Serum ALT levels in 29 week old non-fasting mice. ALT, alanine transaminase. n=11 wild type and n=13 *Slc16a11* knockout. (G) Plasma thyroid hormone levels in 19 week old non-fasting mice. T4, thyroxine; TSH, thyroid-stimulating hormone. n=13 wild type and n=15 *Slc16a11* knockout. (H-I) H&E staining of tissues from 54 week old fasting mice. (H) Images from liver were taken using 20x (200x; scale bar = 50 μM) and 63x (630x; scale bar = 20 μM) objectives. Black boxes indicate regions shown at 63x. The yellow box in *Slc16a11* knockout highlights region with additional immune cell infiltration. (I) Images from subcutaneous and visceral adipose and skeletal muscle were taken using a 20x objective (200x; scale bar = 50 μM) and pancreatic islets were taken using a 40x objective (400x; scale bar = 50 μM).

**Figure S3. Related to Figures 4-5.**
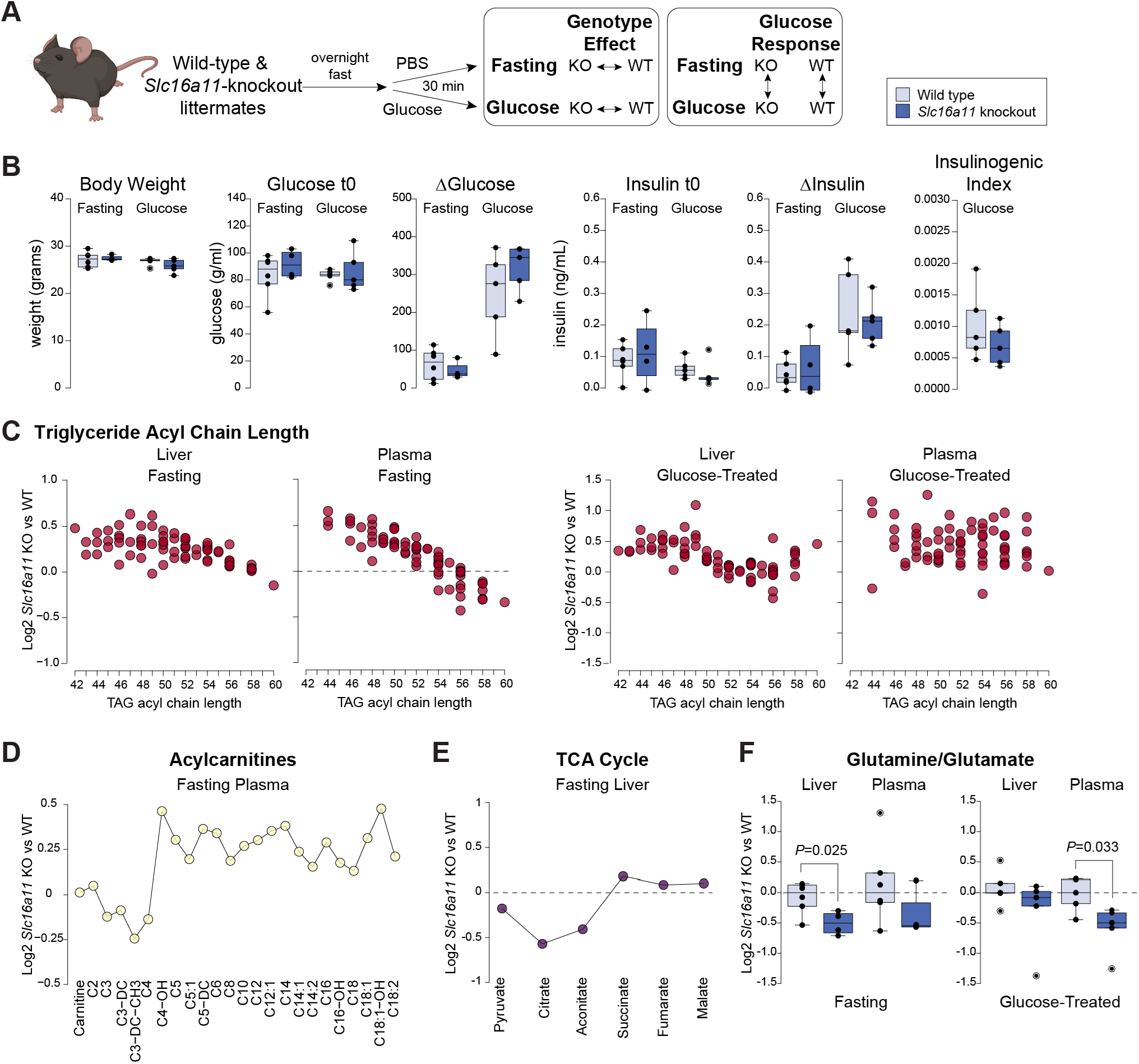
Impact of Mct11 deficiency on metabolite profiles at fasting and following glucose treatment. (A) Depiction of experimental design to assess the *in vivo* impact of Mct11 deficiency on hepatic metabolism after fasting and in response to glucose. (B) Measurements of body weight, glucose, and insulin in 17 week old mice used for metabolomics and RNA-sequencing analyses. Fasting glucose or insulin=t0; change (Δ) in glucose or insulin 30 minutes after glucose administration=t30-t0; Insulinogenic index=[Insulin (t30-t0)/Glucose (t30-t0)]. Fasting mice: n=6 wild type and n=4 *Slc16a11* knockout. Glucose-treated mice: n=5 wild type and n=5 *Slc16a11* knockout. (C) Liver and plasma TAG profiles in fasting and glucose-treated *Slc16a11*-knockout mice. Individual TAGs are grouped by acyl chain length. (D) Plasma acylcarnitine profile in fasting *Slc16a11*-knockout mice. (E) Effect of *Slc16a11* deficiency on hepatic TCA cycle metabolites in fasting mice. (F) Ratio of glutamine-to-glutamate in liver and plasma from fasting and glucose-treated mice. Plots show log2 foldchange of metabolites in *Slc16a11*-knockout relative to wild-type mice.

**Figure S4. Related to Figures 4-6.**
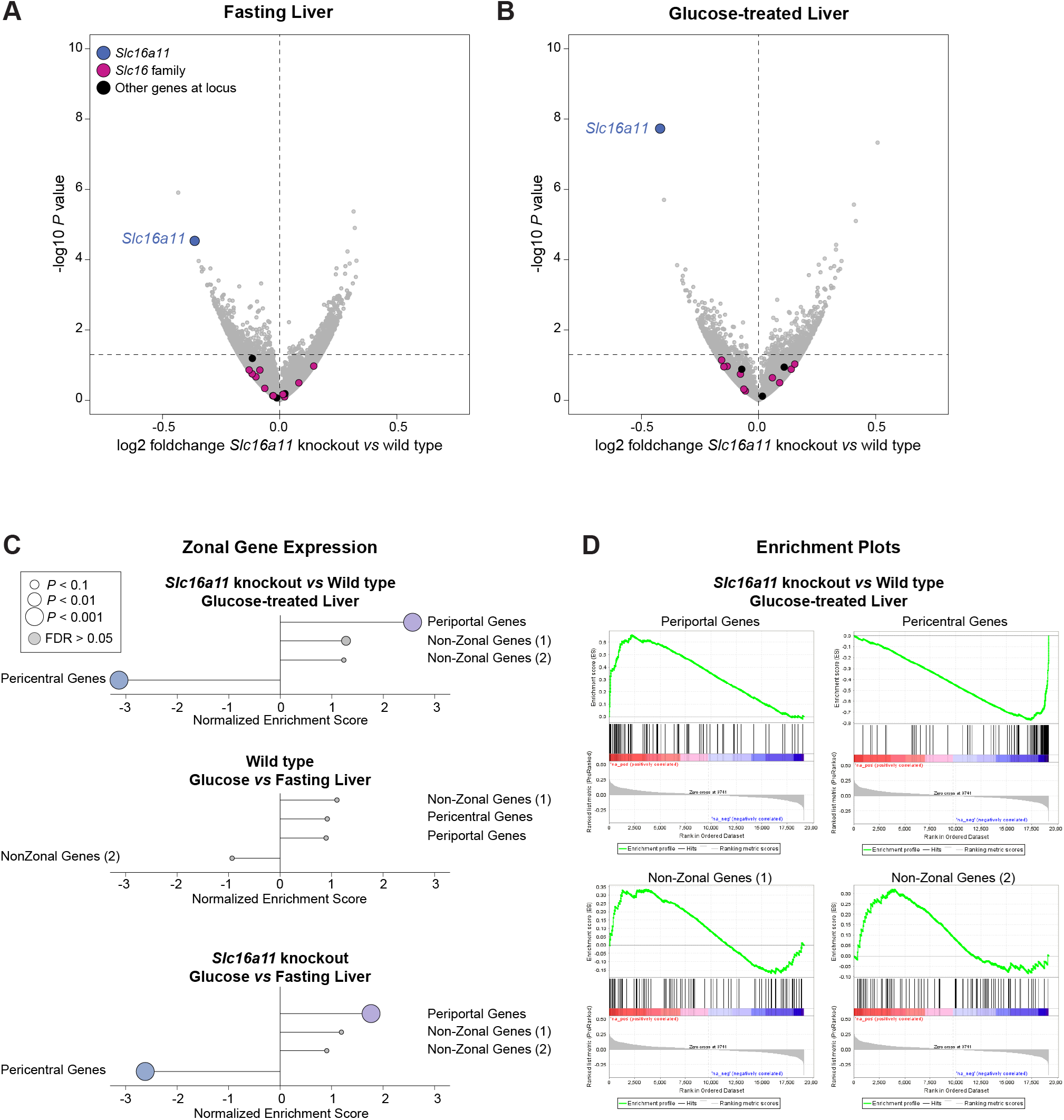
RNA-sequencing analyses of zonally-distributed genes in wild-type and *Slc16a11*-knockout mice. (A and B) Volcano plots showing expression differences in livers from (A) fasting and (B) glucose-treated wild-type and *Slc16a11*-knockout mice. Slc16 family members and other genes within 100 kb of the *Slc16a11* locus (*Bcl6b*, *Clec10a*, *Rnasek*) are indicated. Plots show log2 foldchange in *Slc16a11*-knockout relative to wild-type mice. (C) Gene set enrichment analysis (GSEA) of expression changes in genes with pericentral and periportal zonation profiles. (D) Enrichment plots from GSEA of zonally-distributed and non-zonal genes in glucose-treated *Slc16a11*-knockout versus wild type livers.

**Figure S5. Related to Figures 6.**
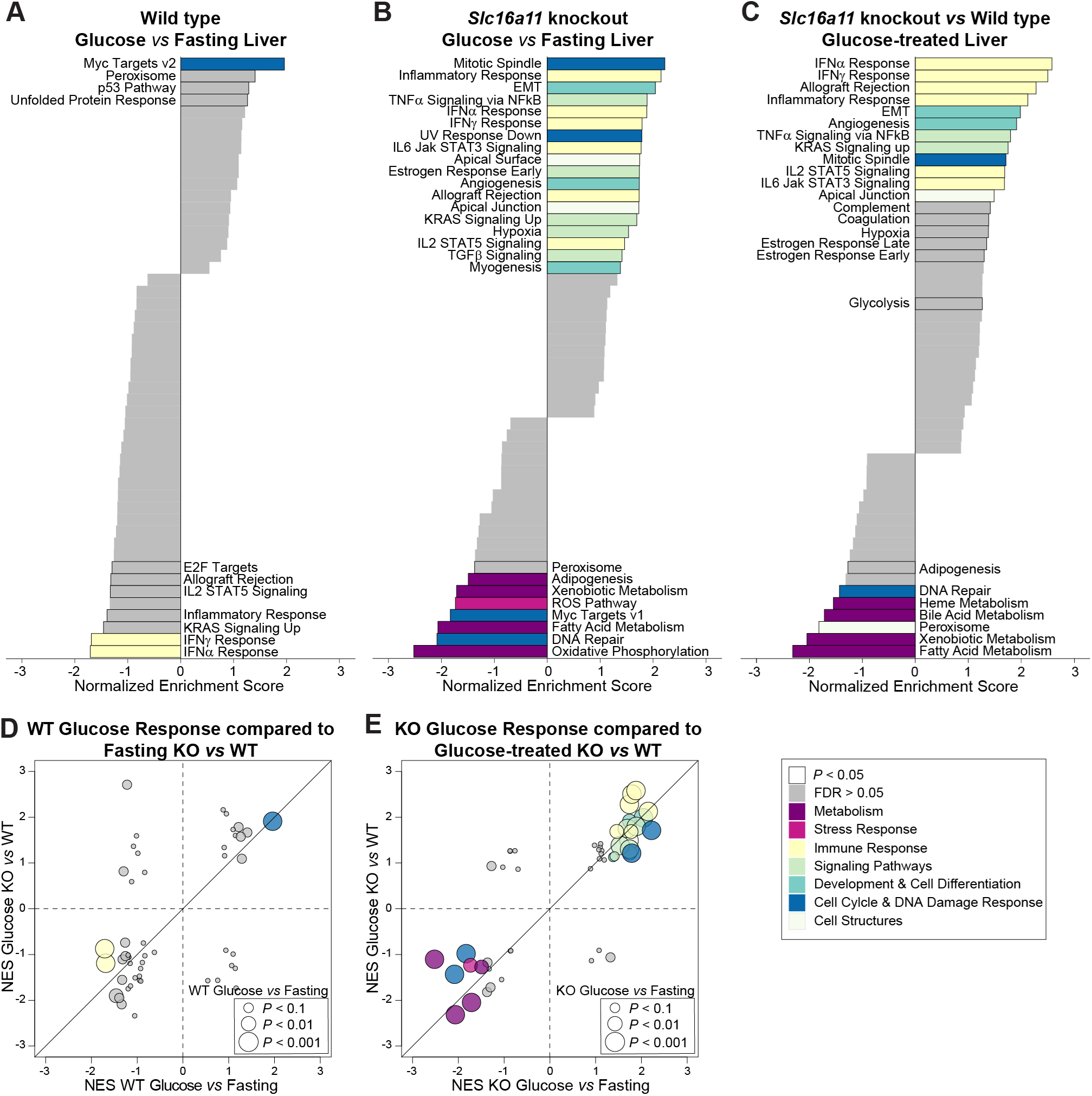
Pathway analyses of gene expression changes in livers from glucose-treated wild-type and *Slc16a11*-knockout mice. (A-C) GSEA of liver transcriptomic profiles comparing (A) glucose-treated and fasting wild-type mice, (B) glucose-treated and fasting *Slc16a11-*knockout mice, and (C) glucose-treated *Slc16a11*-knockout and wild-type littermates. Each bar represents a different pathway from the Hallmark pathways collection (Liberzon et al., 2015). Pathways with *P* <0.05 are labeled and outlined. Colors represent different categories of pathways. Non-significant pathways with (false discovery rate [FDR] > 0.05) are shown in gray. (D) Scatterplot showing transcriptomic pathway changes observed when comparing livers from glucose-treated and fasting wild-type mice with those observed when comparing fasting *Slc16a11*-knockout *vs* wild-type littermates. *P* values and FDR indicate results from comparison of glucose-treated *vs* fasting wild-type mice. (E) Scatterplot showing transcriptomic pathway changes observed when comparing livers from glucose-treated and fasting *Slc16a11*-knockout mice with those observed when comparing glucose-treated *Slc16a11*-knockout and wild-type littermates. *P* values and FDR indicate results from comparison of glucose-treated *vs* fasting *Slc16a11*-knockout mice.

**Figure S6. Related to Figure 6.**
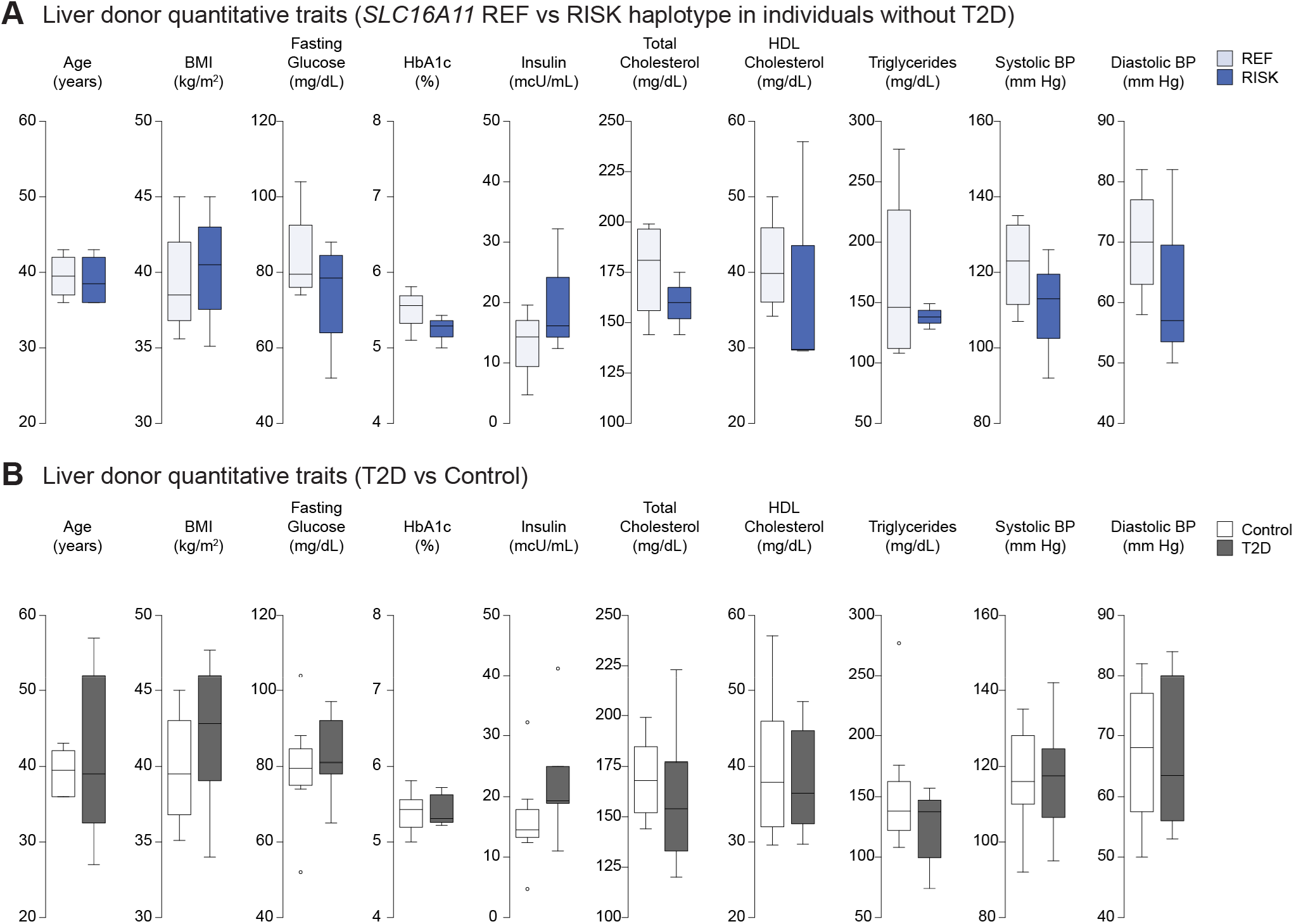
Quantitative traits of human liver donors used in RNA-sequencing analyses. (A and B) Box plots depict quantitative trait analyses in donors used for RNA-sequencing analyses comparing (A) individuals without T2D who are homozygous reference (REF) and homozygous T2D risk (RISK) carriers of the *SLC16A11* haplotype (n=4 each) and (B) individuals with and without T2D (n=8 each). No statistical differences in quantitative traits were detected as assessed using analysis of variance followed by Tukey’s HSD post-hoc test.

## SUPPLEMENTAL TABLE LEGENDS

**Table S1. Related to Figures 2 and S1-S2. Individual metabolite changes in primary human hepatocytes following knockdown of *SLC16A11***

**Table S2. Related to Figures 2 and S1-S2. Metabolic pathway changes in primary human hepatocytes following knockdown of *SLC16A11***

**Table S3. Related to Figures 4-5 and S4. Individual metabolite changes in *Slc16a11-*knockout mice**

**Table S4. Related to Figures 4-5. Metabolic pathway changes in S*lc16a11-*knockout mice**

**Table S5. Related to Figures 4 and 6. Transcriptomic pathway changes in *Slc16a11*-knockout mouse liver**

**Table S6. Related to Figure 6. Transcriptomic pathway changes in human liver from *SLC16A11* T2D risk haplotype carriers and individuals with T2D**

